# AlphaInterp: Mechanistic Interpretability of AlphaFold 3 Reveals How Evolutionary Information Shapes Protein Structure Prediction

**DOI:** 10.64898/2026.04.22.720175

**Authors:** Jonathan Feldman, Jeffrey Skolnick

## Abstract

AlphaFold 3 predicts biomolecular structures with unprecedented accuracy, yet the computations transforming sequence and evolutionary data into structural coordinates remain poorly understood. Here, we present a systematic mechanistic interpretability analysis of AlphaFold 3, tracking its internal representations across the forward pass. Probing four critical network checkpoints reveals that the Pairformer compresses diffuse co-evolutionary inputs into a compact latent geometry where complex biophysical features become linearly decodable. Using causal activation patching, we demonstrate that predicted confidence is directly manipulable within this latent space, allowing geometric certainty to be transferred across entirely unrelated proteins. Furthermore, across adversarial-mutation, fold-switching, and generalization benchmarks, we show that AlphaFold 3’s representational coherence strictly requires comparative evolutionary context. The latent space collapses when multiple sequence alignments are removed, regardless of sequence familiarity or training-set membership. This stability requires phylogenetic diversity rather than alignment depth, and a minimal set of highly divergent homologs is sufficient to anchor the latent space and activate the model’s structural priors. These findings indicate that AlphaFold 3’s representational coherence is deeply tied to evolutionary scaffolding, suggesting it functions similarly to an advanced fold-recognition system and highlighting that protein structure prediction from sequence alone is not yet fully solved.

## 1 Introduction

The release of AlphaFold 2 in 2021 fundamentally transformed structural biology [1, 2]. By achieving near-experimental accuracy on a substantial fraction of protein targets, it demonstrated that deep learning could solve what had been considered one of biology’s grand challenges: predicting three-dimensional protein structure from amino acid sequence [3]. AlphaFold 3, released in 2024, extended these capabilities to nucleic acids, small molecules, and complete biomolecular assemblies, with improved accuracy for protein-protein interfaces and multimeric complexes [4]. As a result, AlphaFold 3 has rapidly become the de facto standard in computational structural biology, underpinning an expanding ecosystem of derivative tools spanning protein design, binding affinity prediction, fitness modeling, and experimental structure determination [5–15].

The pervasive integration of AlphaFold into research workflows makes understanding its underlying capabilities—and limitations—a matter of broad scientific importance [16–20]. If AlphaFold has learned the true biophysical principles relating sequence to structure, then the tools built upon it inherit a robust foundation. If its predictions are instead primarily driven by pattern matching to training examples, derivative methods may be inheriting the same constraints, effectively tethered to the training distribution regardless of the conformation a given sequence will actually adopt. The distinction is not trivial: models without genuine biophysical grounding are unlikely to generalize to novel protein families, engineered sequences, or proteins with substantial mutations relative to characterized homologs [21, 22].

There is mounting evidence that current structure prediction models struggle with such out-of-distribution inputs. AlphaFold’s performance degrades on proteins with low sequence similarity to its training set [12, 21, 23, 24]. Its predictions for alternative conformations and fold-switching proteins often converge to a single dominant fold rather than reflecting known structural diversity [10, 21, 23]. When used for protein design, AlphaFold-based pipelines can generate sequences that appear well-folded in silico but fail experimentally [9, 25, 26]. A particularly striking manifestation of this brittleness is mutational invariance: predicted structures can remain nearly unchanged under aggressively deleterious point mutations, tolerating mutational loads that standard biophysics would predict to be highly destabilizing [21]. This phenomenon has been documented at the level of predicted coordinates, but its representational origins—whether the invariance is embedded in the model’s internal geometry or emerges only at the final structure module—have not been examined.

Addressing these questions requires going beyond standard benchmarks, which measure predictive accuracy but do not test whether models capture causal sequence-structure relationships. It requires opening the model itself. Mechanistic interpretability— the systematic analysis of what information is encoded in a model’s internal representations, how that information is organized, and whether it is causally upstream of model outputs—has proven powerful for understanding large language models and biological sequence models such as ESM [27–29], but has not yet been applied systematically to structure prediction models like AlphaFold 3. The internal representations of AlphaFold 3—its single residue-level channels and its pairwise interaction channels—carry the information that determines the predicted fold, but what exactly they encode, how they evolve through the network, and what role the multiple sequence alignment (MSA) plays in shaping them has remained opaque [4].

Here, we develop a mechanistic interpretability framework for AlphaFold 3 and apply it across a suite of carefully designed benchmarks. Extracting single and pair representations at four checkpoints spanning the full forward pass, we ask: what biophysical features are linearly encoded in these representations, and when do they become accessible? How do the pair and single channels differ in their informational content? Is structural uncertainty encoded in the internal geometry, and can it be causally manipulated? How does mutational invariance manifest at the representational level, and from where does it originate? How do the representations respond to structurally similar but sequentially dissimilar proteins and to sequentially similar but structurally divergent ones? And critically, how does the MSA shape the internal representations—and what happens to those representations when it is removed?

The answers to these questions paint a coherent and, in some respects, surprising picture of how AlphaFold 3 constructs its predictions, where its power originates, and where its fundamental limitations lie.

## 2 Results

### 2.1 Checkpoint Extraction and Experimental Design

To move beyond AlphaFold 3’s outputs and begin to unwrap its internal logic, we extracted its single (*L* × 384) and pair (*L* × *L* × 128) representations at four checkpoints spanning the full forward pass (see Methods), each corresponding to a distinct computational phase in the transformation of sequence information into structural geometry [4]. Checkpoint A is sampled immediately after sequence initialization but before the MSA module has been fully applied. It is not entirely MSA-free: per-position amino acid frequency distributions and deletion means are already concatenated into the initial single representation during input preparation, embedding a shallow statistical summary of co-evolutionary context before any deep pairwise interaction occurs. Checkpoint B is sampled after the MSA module and before the Pairformer, collected exclusively at the first recycling iteration (*r* = 0). This marks the first point at which deep co-evolutionary information enters the pair channel through the outer product mean and row-wise gated self-attention. Checkpoint C_1_ is sampled after the final Pair-former layer at *r* = 0, capturing the model’s first complete bottom-up structural hypothesis from sequence and MSA alone, with no geometric prior carried in from previous recycling iterations. Checkpoint C_N_ is sampled after the final Pairformer layer at the terminal recycling iteration *r* = *N* − 1 [4]. It is the representation that directly conditions the diffusion module and determines the predicted structure. For all analyses, *N* = 10, consistent with the AlphaFold 3 default [4]. Additional details on checkpoint selection and extraction are provided in Methods.

To determine what these internal states encode, we assembled a dataset of 400 monomeric proteins: 200 with less than 30% sequence homology to any protein plausibly present in the AlphaFold 3 training set (Novel), and 200 carrying no such restriction (Similar). Crucially, all proteins were deposited after September 30, 2021, the AlphaFold 3 training cutoff, ensuring that even the Similar set is temporally out-of-distribution [4, 5]. This design allows us to probe generalization and potential memorization separately throughout the analyses that follow. Because embedding dimensionality scales with protein length, direct comparison across proteins requires reduction to fixed-size vectors. We systematically evaluated multiple pooling strategies (see Methods) and found that simple mean pooling was optimal for global prediction tasks. Single embeddings were mean-pooled along the residue dimension to yield 384-dimensional vectors, whereas pair embeddings were reduced via upper-triangular mean pooling to yield 128-dimensional vectors (see Methods).

### 2.2 The Pairformer Reorganizes the Embedding Space Around Structural Quality

We begin with a simple but direct baseline question: can AlphaFold 3’s internal representations predict the confidence of the structures they themselves produce? This form of self-prediction provides a measure of representational self-consistency—the extent to which the latent geometry at a given checkpoint anticipates the model’s final structural judgment. Using ridge regression, which performed comparably to other tested regressors (see Methods), we find a striking progression [30, 31]. For pair representations, the coefficient of determination (*R*^2^; see Methods) for predicting the AlphaFold 3-predicted Template Modeling score (pTM) [4], a measure of global structural confidence, increases from 0.34 at Checkpoint A to 0.86 at Checkpoint C_N_. Across all stages, pair representations consistently outperform single representations, as can be seen throughout Figure 1.

**Fig. 1.**
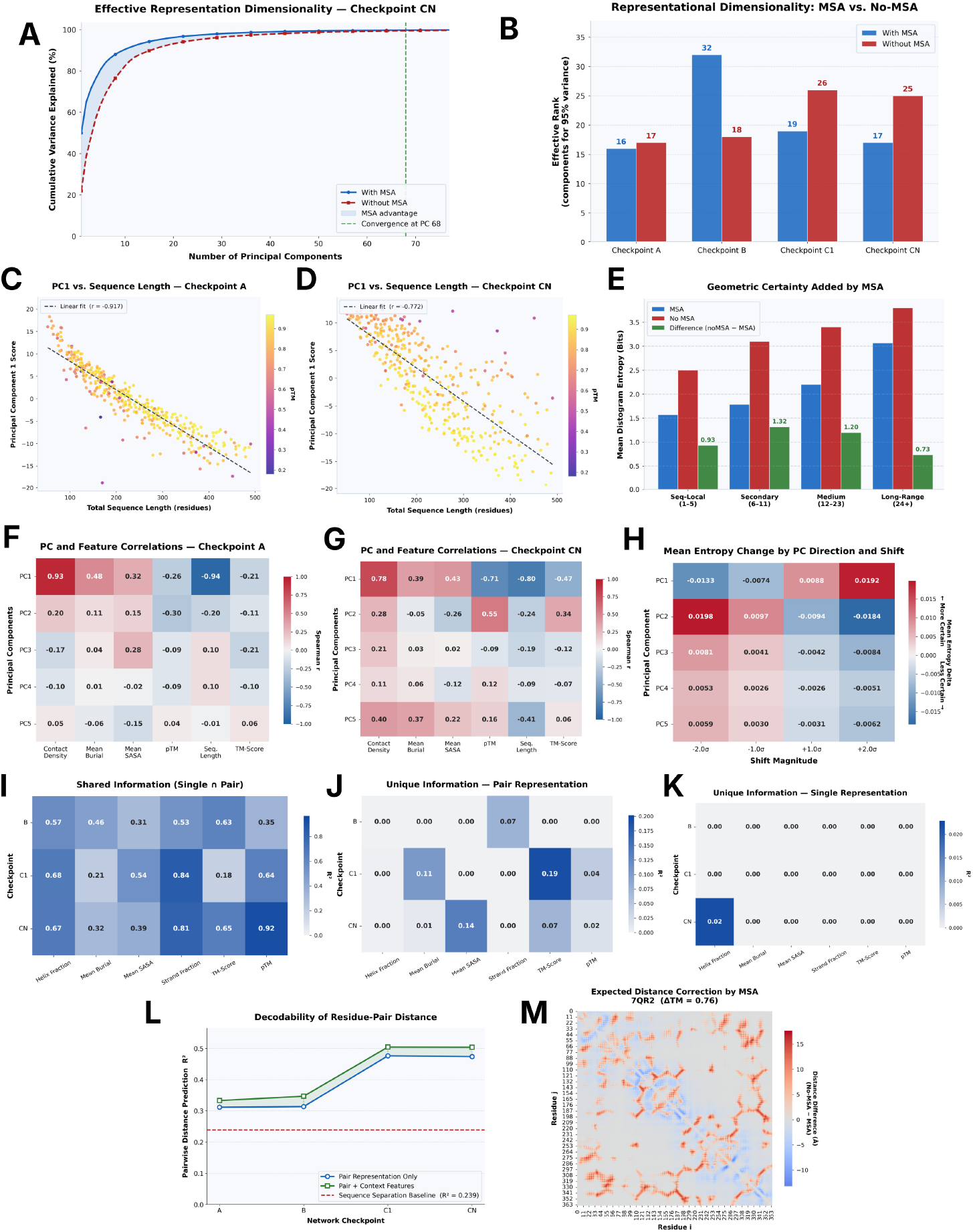
AlphaFold 3 internal checkpoints and representational self-consistency. **(A)** Cumulative explained variance vs. Principal Components (PCs)—orthogonal directions capturing maximal variance in the data—for the *C*_*N*_ pair representation (blue: MSA; red: no-MSA); the MSA curve saturates faster, indicating a more consolidated space. **(B)** PCs required for 95% variance across all four checkpoints; checkpoint *B*’s expansion is attributable to MSA integration. **(C–D)** PC1 score vs. sequence length at *A* and *C*_*N*_ (colored by pTM); the association, as measured by the Pearson correlation coefficient, weakens at *C*_*N*_, implying richer structural encoding at later layers. **(E)** Mean distogram entropy for MSA, no-MSA, and their difference by sequence separation bin; the gap peaks at intermediate separations. **(F–G)** Spearman correlations between the top five pair-representation PCs and biophysical features at *A* and *C*_*N*_ ; predictive power increases with network depth. **(H)** Distogram entropy change as PCs are shifted by varying magnitudes, linking representational geometry to model certainty. **(I–K)** Shared (I), unique-pair (J), and unique-single (K) information across *B, C*_1_, *C*_*N*_ ; pair representations encode more unique structural information. **(L)** Distance decodability (*R*^2^) across all checkpoints for a sequence-separation baseline, an isolated pair probe, and a context-augmented probe. See Methods for more details. **(M)** Expected pairwise distance difference (no-MSA − MSA) for 7QR2; positive values indicate pairs where no-MSA over-estimates separation. The ΔTM-score is 0.76 in the MSA-enriched model’s favor.

Part of this result is mechanistically expected: the final pair representations feed directly into both the distogram head and the confidence module, making them immediate informational determinants of both structure and confidence [4]. The more interesting question is where the gain occurs. The largest transformation is not explained by MSA injection alone—it is driven by the Pairformer itself. Even after the first complete bottom-up pass at C_1_, recycling continues to reshape the latent geometry, progressively aligning it with structural confidence. The Pairformer seems not to merely refine the MSA-derived representation—it reorganizes the entire embedding manifold around the notion of fold quality. That this geometric sharpening continues through late recycling suggests that confidence is not a static change made at a single stage but an emergent property of the iteratively evolving latent space.

The principal component analysis (PCA) of the pooled representations reveals the mechanism underlying this progression [32]. The effective dimensionality of the pair representations—defined as the number of principal components required to explain 95% of the variance—largely follows a trajectory of compression: A(16) → B(32) → C_1_(19) → C_N_(17) (see Figures 1A and 1B), where the numbers in parentheses are the number of principal components. The transient expansion at Checkpoint B can be interpreted as the MSA module injecting co-evolutionary information broadly across the latent space, simultaneously activating many directions of variance. What follows is more revealing. As the Pairformer and recycling layers operate, this high-dimensional co-evolutionary signal is compressed into a far more compact and geometrically coherent subspace. The compression mirrors almost exactly the rise in self-consistency observed in the pTM probes, suggesting that the Pairformer’s central function is to distill co-evolutionary statistics and geometrical knowledge into a concentrated representational and, subsequently, structural manifold.

The evolution of the leading principal component makes this transformation apparent. At Checkpoint A, PC1 is almost perfectly negatively correlated with sequence length (Pearson *r* = −0.917 and Spearman *ρ* = −0.943; Figures 1C and 1F), meaning the dominant axis of variation is a trivial feature with little structural content. By Checkpoint C_N_, that axis has been partially repurposed. The correlation with sequence length diminishes (Pearson *r* = −0.77 and Spearman *ρ* = −0.80; see Figures 1D and 1G). Simultaneously, PC1 begins to strongly correlate with pTM (Spearman *ρ* = −0.71; see Figures 1F and 1G). The latent manifold has not merely been refined— it has been shifted to more strongly reflect structural confidence. As we show below, this reorientation extends to other biophysical properties as well.

Whether this geometric reorganization is merely descriptive or whether it actively drives downstream predictions is the key question separating correlation from causality. To test this, we performed activation patching directly on the late-stage pair representations at Checkpoint C_N_ (see Methods), intervening on the internal geometry before the distogram head and measuring the downstream consequence [4, 33].

The dominant principal components prove to be causally active. Shifting a representation by +1*σ* along PC1 increases distogram entropy by only +0.0044 for highly accurate predictions but produces a +0.0133 entropy increase for poorly predicted targets (an overall average of +0.0088, as can be seen in Figure 1H)—a three-fold amplification that strongly inversely correlates with the model’s original predicted confidence (Spearman’s *ρ* = −0.757). The sensitivity of the downstream structural distribution therefore depends directly on where the representation sits within the latent confidence manifold: accurate predictions occupy a geometrically stable region, while poor predictions are fragile. This geometric control extends beyond PC1, as shifting representations along PC2 uniformly reduces distogram entropy by −0.0198, indicating that multiple principal directions actively shape the model’s geometric certainty (Figure 1H; see Methods for more details).

Cross-protein activation patching sharpens the result further. Transplanting the *C*_*N*_ representation from a near-perfect structural prediction (TM-score = 0.999, where TM-score measures global structural alignment quality on a scale of 0 to 1 [34]) into poorly predicted targets compels the downstream distogram head to emit highly certain, tightly packed contact maps, dropping mean distogram entropy by −0.268 across those targets. The internal geometry encoding a successful fold can therefore be transferred between unrelated proteins and still impose structural certainty on the downstream distribution. For more details, refer to the Methods section.

### 2.3 Biophysical Features Are Linearly Encoded—Primarily in the Pair Representation

Having established how the geometry of the latent space evolves, we next asked what, exactly, is encoded within it, and when does that information become accessible. To answer this, we applied linear probes to representations extracted at each checkpoint. For each of the 400 proteins, we extracted per-residue secondary structure, solvent-accessible surface area (SASA), residue burial depth, C*α*–C*α* contact maps within 8 Å, and residue-pair Euclidean distance from PDB coordinates. Probes were applied to single representations for per-residue scalar properties and to pair representations for pairwise properties (see Methods).

A consistent pattern emerges across all features, where predictive accuracy improves progressively through the pipeline, indicating that biophysical signal is continuously refined rather than abruptly introduced at any single stage.

As can be seen in Figure 1L, for pairwise *C*_*α*_–*C*_*α*_ Euclidean distance, *R*^2^ increases from 0.311 at Checkpoint A to 0.474 at Checkpoint C_N_ using only the paired representations, and to 0.504 using local context features (See Methods). *C*_*α*_–*C*_*α*_ contact prediction follows the same trajectory, with balanced accuracy rising from 0.642 to 0.703. For secondary structure, probed from AlphaFold 3’s per-residue single representation—where each residue is assigned its own 384-dimensional embedding vector—accuracy improves from 0.43 at Checkpoint B to 0.57 at Checkpoint C_N_, while SASA (0.02 → 0.09) and burial depth (0.05 → 0.15) show smaller but directionally consistent gains. As a necessary baseline, AlphaFold 3’s per-residue single representations at all checkpoints achieve near-perfect accuracy in amino acid identity reidentification, confirming that residue identity is preserved throughout the forward pass and that the probes are operating on a faithful encoding.

The modest absolute values for some features, particularly SASA, are not unexpected. There is no a priori reason to expect AlphaFold 3 to organize these properties into linearly separable axes—it is entirely plausible that they exist as non-linear superpositions within the representation that a linear probe cannot recover. What is notable is not the ceiling of performance but its consistency: residue contacts, pairwise distances, and secondary structure all become increasingly linearly decodable—that is, predictable by a linear machine learning probe—as the model progresses, indicating that these structural properties are not merely present in the representations but are organized in a progressively more accessible geometric form.

Critically, this signal is not evenly distributed across the two channels. To quantify the asymmetry, we performed an *R*^2^ shared-information decomposition for each predictive target (see Methods; Figures 1I-K), where *R*^2^ is the coefficient of determination. Across pTM, TM-score, SASA, burial depth, and secondary structure, the single and pair representations share a substantial fraction of variance, reflecting their joint participation in the forward pass [4, 34]. Yet the pair representation consistently retains greater independent predictive power, establishing that the structural signal is not simply duplicated across channels but is concentrated in the pair representation.

These results delineate a clear functional specialization within AlphaFold 3. The single representation acts as a residue-level state tracker, encoding sequence composition, length, and global confidence, but remains comparatively insensitive to the local backbone geometry that determines fold. The pair representation carries the burden of geometry, as it encodes relational structure between residues and is progressively sculpted, through the MSA module and Pairformer, into a linearly accessible representation of three-dimensional protein structure. It is this pair manifold, not the single channel, that constitutes the primary geometric substrate of AlphaFold 3’s structural reasoning.

### 2.4 MSA Information Increases Accuracy and Reduces Representational Uncertainty

The analyses above were performed under AlphaFold 3’s standard operating regime, with full MSA information available for every target. The natural next question is what happens when that evolutionary context is removed entirely. To isolate the role of co-evolutionary signal in shaping the latent space, we reran the same 400 proteins without MSAs—a perturbation that is mechanistically revealing in its own right and lays the foundation for the more systematic analyses of MSA depth and fold-switching that follow.

The structural consequences are immediate and severe. Mean TM-score collapses from 0.9363 (std = 0.0987) when MSAs are used to 0.5388 (std = 0.2333) without them. Yet the representational story is more nuanced. Even without MSA information, the no-MSA embeddings retain strong self-predictive power for pTM (*R*^2^ = 0.822) and TM-score (*R*^2^ = 0.845), with accuracy continuing to improve across checkpoints. By contrast, the MSA-conditioned embeddings show low self-predictive power for these same quantities—not due to representational failure, but as an artifact of limited dynamic range. MSA-conditioned predictions are compressed into a narrow, high-confidence distribution whose low variance renders regression effectively uninformative, regardless of the underlying signal. The no-MSA condition, by exposing substantially greater output variance, allows that signal to become linearly measurable. Both regimes track structural quality internally, but only one makes that tracking visible to a probe. Additionally, as shown in Figures 1A and 1B, the effective dimensionality of the paired representations in the no-MSA condition at later checkpoints—C_1_ and C_N_—is substantially higher, indicating a corresponding decrease in representational homogeneity in the absence of MSA information.

Another informative result emerges from comparing the distograms generated from Checkpoint C_N_ pair representations with and without MSA for identical input sequences [35]. As can be seen in Figure 1E, MSA removal globally increases distogram Shannon entropy, reflecting a broad rise in uncertainty over inter-residue distances. This uncertainty is not uniformly distributed across the contact map, with the entropy increase being maximal for residue pairs separated by 6–23 sequence positions (Figure 1E), while still being substantially high at other ranges, indicating that the MSAs most bolster model confidence—though not structure (see MSA Perturbation section)—at local secondary-structure contacts rather than long-range tertiary interactions.

KL-divergence between MSA and no-MSA distograms reinforces this interpretation quantitatively [36]. KL-divergence correlates strongly with the corresponding change in TM-score relative to the PDB ground truth (Pearson *r* ≈ 0.72 across all sequence-separation bins), directly linking the information-theoretic displacement of the model’s internal distance distribution to realized structural collapse in 3D space. KL-divergence thus serves as a mechanistic bridge between latent uncertainty and downstream structural failure.

Finally, we separated the Similar and Novel cohorts to test whether training-set membership modulates any of these effects. Across all biophysical probes— secondary structure, SASA, burial depth, contact prediction, and pairwise distance— performance differences between the two groups were negligible at every checkpoint, despite Similar proteins being supported by MSAs approximately twice as deep (mean depth 9,513 vs. 4,587). AlphaFold 3 therefore encodes biophysical information in a manner that is largely invariant to training-set familiarity. That MSA depth differences of this magnitude leave the representations unchanged is itself informative and suggests that the decisive factor is not the quantity of co-evolutionary signal but its presence or absence—a hypothesis we test directly in the sections that follow.

### 2.5 Mutational Invariance Is a Representational Phenomenon, Not Just a Structural One

We next turned to a recently noted and consequential behavior in AlphaFold-like systems: mutational invariance. Prior work has shown that predicted structures can remain strikingly stable under aggressively deleterious sequence perturbations, tolerating large mutational loads before collapsing [21]. This phenomenon has generally been characterized as a structural observation, measured at the level of final predicted coordinates. Here, we asked whether that robustness is merely an emergent property of the output structure or whether it is already embedded within the model’s internal representations.

All 400 proteins were subjected to an adversarial mutational analysis following the protocol of [21]. Residues were replaced with purposefully deleterious point mutations at loads of 10, 20, 40, 70, and 80%, where mutations were cumulative as percentages increased for a given protein, and the resulting structures were compared to the original unmutated predictions. As previously reported, predicted structures exhibit striking invariance up to 40% of their sequence being deleteriously mutated, after which they collapse. We extended this perturbation directly into the representational interior of the model, extracting embeddings at Checkpoints A, B, C_1_, and C_N_ for every mutated sequence. Because we are comparing the same proteins across mutational thresholds, no pooling was applied—the residue-length dimension remains identical across conditions (see Methods).

The internal representations mirror the structural behavior with remarkable fidelity (Figures 2A and 2E). Cosine distance between original and mutated representations increases in lockstep with TM-score divergence between mutated and unmutated predicted structures, crossing the threshold of major change between 40% and 70% mutation, where it reaches approximately 0.731. This correspondence is quantitatively tight, where cosine distance is a strong predictor of TM-score divergence, with a Pearson correlation coefficient of −0.948 and a Spearman correlation coefficient of −0.899, as shown in Figure 2C. Mutational invariance is therefore not only a property of the final fold but of the latent space itself. The model’s resistance to sequence perturbation is not an artifact of the diffusion module or any single downstream subsection but a deep-rooted property of the representational pipeline—the same threshold behavior visible in the predicted coordinates is already present in the embeddings, indicating that a stable internal structural hypothesis is maintained long before coordinates are emitted.

**Fig. 2.**
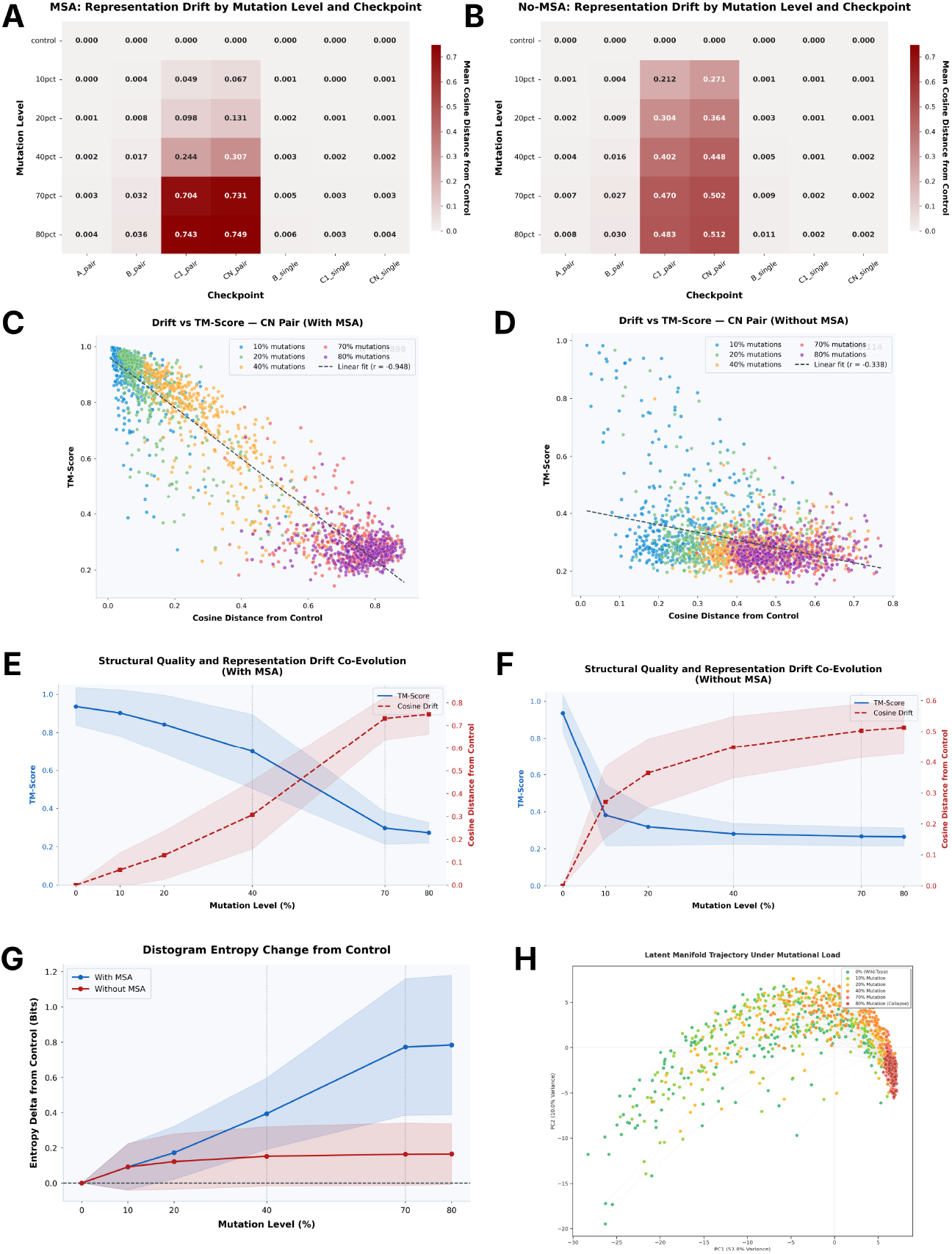
Mutational robustness of AlphaFold 3 internal representations. **(A–B)** Mean cosine distance of MSA-enriched (A) and no-MSA (B) pair and single representations from their respective unmutated controls, shown across all mutation levels. **(C–D)** Scatterplots of *C*_*N*_ pair embedding cosine distance from control vs. TM-score, for MSA-enriched (C) and no-MSA (D) conditions, colored by mutation level. The relationship between representational drift and structural degradation is substantially more linear and monotone under MSA conditioning. **(E–F)** Dual-axis line plots showing TM-score (left axis) and cosine distance from the unmutated control at C_N_ (right axis) across mutation levels, for MSA-enriched (E) and no-MSA (F) conditions, with shading indicating one standard deviation. Under MSA conditioning, representational drift tracks structural degradation closely; without MSA, drift saturates rapidly at high levels while TM-score continues to fall, indicating a decoupling of representational and structural change. **(G)** Mean distogram entropy across mutation levels for MSA-enriched and no-MSA representations. The no-MSA baseline entropy is elevated throughout and increases less steeply with mutation, suggesting the model operates under persistently higher geometric uncertainty that is less responsive to mutational load. **(H)** PCA projection of pooled MSA-conditioned *C*_*N*_ pair representations, colored by mutation level. Under increasing mutational load, embeddings converge toward a restricted region of the representational manifold, revealing a shared geometric trajectory that characterizes the model’s transition into a regime of structural uncertainty.

The localization of this signal is especially revealing. Despite extreme sequence modification, representational changes are confined almost entirely to the pair representations at Checkpoints C_1_ and C_N_ (Figure 2A). No meaningful drift is observed at Checkpoints A or B. The uncertainty introduced by mutation is not expressed during sequence initialization or during the immediate integration of MSA information—it emerges specifically within the Pairformer, where residue-residue geometry is actively reconciled into a fold hypothesis. Even severe sequence corruption is absorbed by the earlier stages of the model; only when the Pairformer attempts to impose geometric consistency does the latent space begin to diverge.

We next asked whether the resulting collapse follows protein-specific trajectories or converges onto a shared failure geometry. The answer is unambiguous. As mutation load approaches 70–80%, the representational drift vectors of individual proteins, projected into PCA space, converge onto a nearly identical direction—a mean resultant vector length of 0.901 out of a maximum of 1.0 (see Methods)—indicating that structural collapse is not protein-specific but follows a shared geometric trajectory (Figure 2H). PC1 serves as the dominant manifold for this degradation, absorbing the vast majority of the representational shift (mean absolute component = 0.767), with subsequent axes capturing progressively less of the failure signal (PC2 = 0.325, PC3 = 0.239). Projected into the subspace of these top three components, the drift vector points reliably toward a shared, fold-invariant region of the latent space. The model does not fail randomly under extreme mutation; instead, it collapses along a universal geometric pathway.

This interpretation is supported by the observed dependence of collapse magnitude on original prediction quality. Proteins that were already poorly predicted prior to mutation—those with an initial TM-score below 0.5 relative to the PDB reference structure—shift by about 0.548 in cosine distance at 80% mutation. In contrast, proteins with an initial TM-score of ≥ 0.9 diverge by 0.763 under the same perturbation. This suggests that proteins for which the model was already uncertain tend to exhibit smaller additional shifts, consistent with their representations already lying closer to a collapsed regime. Correspondingly, larger geometric changes are more often observed when the initial latent state is more structured. While this pattern is consistent with a collapse-like behavior under mutation, it is more conservatively interpreted as indicating that the model’s representations become progressively less discriminative as sequence perturbations increase, rather than definitively establishing a discrete failure mode.

Examining which proteins most strongly resist this collapse reveals clear structural correlates. Longer proteins are substantially more resistant (Spearman *ρ* = −0.516), likely because larger chains distribute the mutational burden across more residues, requiring a greater fraction of the sequence to be overcome before the model’s geometric expectations are destabilized. Proteins that collapse earliest exhibit higher average SASA (Spearman *ρ* = 0.338) and higher contact density (Spearman *ρ* = 0.501), identifying small, tightly packed, solvent-exposed structures as the most susceptible class. These proteins likely depend on a concentrated set of local geometric constraints, making their latent fold hypothesis easier to disrupt once enough key residues are perturbed. Even so, mutation loads exceeding 40% are required to overcome both structural and representational stability even in this most vulnerable class— underscoring that the model’s robustness to deleterious sequence perturbation is embedded deeply within its pair-level geometric reasoning, not a surface property of the final predicted structure.

### 2.6 Mutational Invariance Is MSA-Dependent

The mutational invariance described above could in principle be attributed to the MSA itself. When a sequence is mutated, its MSA naturally contains homologous variation at many of the perturbed positions, buffering the representational impact and stabilizing the downstream fold hypothesis. A previous study [21] found no major structural difference when MSAs were regenerated versus held fixed under sequence mutation, consistent with this interpretation. Our representational analysis adds a critical nuance: mutational drift does not emerge until the Pairformer, meaning the immediate integration of MSA information at Checkpoint B is insufficient to track sequence perturbation. This raises a direct question—does removing the MSA altogether change the model’s mutational sensitivity?

It does, dramatically. Without an MSA, representational changes emerge immediately: cosine distance at Checkpoint C_N_ reaches 0.271 at just 10% mutation, far earlier than in the MSA-conditioned setting (Figures 2B and 2F). Yet the magnitude then plateaus, becoming largely invariant beyond 40% mutation. The model collapses into its failure regime early and has little geometric structure left to lose thereafter. This collapse is expressed simultaneously in the structural outputs: TM-scores are already catastrophically low at 10% mutation, and entropy changes under mutation are far smaller than in the MSA-conditioned model, confirming representational rigidity rather than sensitivity (Figure 2F).

The contrast between conditions is stark. At 10% mutation, no-MSA structures are wholly inaccurate, while MSA-conditioned structures remain largely invariant. At 40% mutation—a load expected to significantly destabilize the native fold—the MSA-conditioned model still maintains a confident, accurate prediction (TM-score ≈ 0.70). The MSA is not merely buffering local sequence changes at the input stage. It is what allows the Pairformer to sustain a coherent geometric hypothesis at all. Without it, the sequence carries insufficient information to anchor structural reasoning, and even modest perturbations are enough to collapse the representational space entirely.

### 2.7 Fold-Level Clustering in Representation Space Requires Co-Evolutionary Context

To probe AlphaFold 3’s internal representations under a deliberately difficult generalization setting, we turned to SABmark, a benchmark of sequentially dissimilar yet structurally similar proteins. We assembled 100 proteins across 5 groups of 20, each group sharing experimentally validated fold-level similarity while every within-group pair maintains less than 25% sequence homology. Of the 950 possible aligned pairs, 560 carried full pairwise structural overlap annotations and were retained for analysis. As in the mutational analysis, full representations were used without pooling, as the residue subsets being compared were matched exactly in size through the provided experimental alignments (see Methods).

Because SABmark includes residue-level structural alignments, we could ask a sharper question than simple structure-level agreement: do AlphaFold 3’s internal representations recapitulate experimentally validated structural correspondence even when sequence similarity is absent? For each aligned protein pair, we compared pair and single embeddings restricted to the experimentally aligned residues. Two controls were constructed to establish null experiments: for each protein pair with *x* aligned residues, a random sample of *x* non-aligned residues from the same proteins served as an internal control, and a random pair of monomers from the 400-protein dataset aligned over the same number of residues served as an external null.

The results show a clear separation in representational space. Cosine similarity between aligned regions differs significantly from both control conditions, with the strongest signal appearing in the pair representations at C_1_ and C_N_ (Figure 3A). Critically, this signal accumulates only after the Pairformer runs. Checkpoints A and B remain comparatively invariant, consistent with the absence of obvious early sequence features that would reveal structural similarity among sequence-dissimilar proteins. It is only after residue-residue geometry is iteratively reconciled in the Pairformer that the latent space recovers the experimentally validated fold overlap—further evidence that the Pairformer is the primary structural conditioner within AlphaFold 3. Furthermore, the distogram entropy of the fold-switching region is, on average, higher than that of the non-fold-switching region, as can be seen in Figure 3E, indicating that the model has greater structural uncertainty for a region with multiple conformations.

**Fig. 3.**
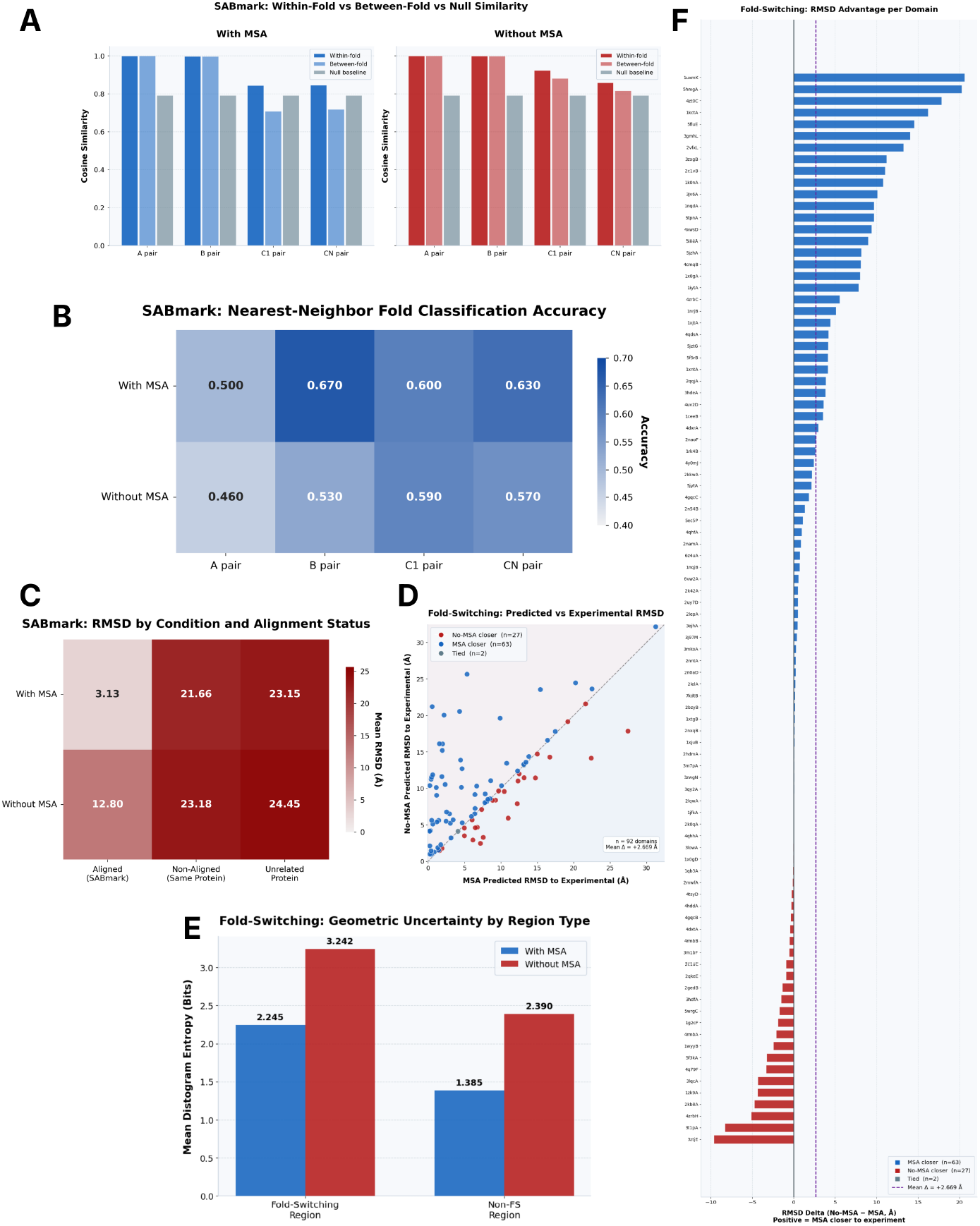
SABmark fold discriminability and fold-switching structural accuracy under MSA conditioning. **(A)** Within-fold, between-fold, and null baseline cosine similarity for all four pair-representation checkpoints (*A, B, C*_1_, *C*_*N*_ ) under MSA-enriched and no-MSA conditions. Without MSA, the within- and between-fold similarities converge at *C*_1_ and *C*_*N*_, indicating that evolutionary context is required to maintain fold-discriminative geometry. **(B)** Heatmap of nearest-neighbor fold classification accuracy for each checkpoint and condition, quantifying how well individual pair representations support recovery of SABmark group membership. **(C)** Mean Kabsch RMSD across 560 SABmark pairs at aligned positions, non-aligned positions within the same proteins, and positions from unrelated proteins in the 400-monomer dataset, for MSA-enriched and no-MSA conditions. MSA conditioning reduces aligned RMSD, while non-aligned and unrelated RMSDs remain comparably high across both conditions. **(D)** Per-domain scatterplot of Kabsch RMSD to the experimental PDB structure at the fold-switching region, comparing MSA-enriched and no-MSA predictions directly. Points above the diagonal indicate domains where the no-MSA structure is less accurate; the majority of domains fall above, showing that MSA conditioning produces systematically closer predictions. **(E)** Mean distogram entropy at fold-switching and non-switching residues under MSA-enriched and no-MSA conditions. Fold-switching residues exhibit higher geometric uncertainty in both conditions, and the absence of MSA further elevates entropy across both region types. **(F)** Per-domain waterfall plot of RMSD delta (no-MSA − MSA) at the fold-switching region, sorted by magnitude. Positive values indicate domains where the MSA-enriched structure is closer to the experimental PDB structure; the MSA-enriched prediction is more accurate in the large majority of cases.

To confirm that this representational signal reflects genuine geometric correspondence rather than an internal artifact, we examined the final predicted coordinates directly. Aligning predicted structures at the experimentally matched residues yields a mean Root Mean Square Deviation (RMSD) of approximately 3.13 Å, whereas non-aligned residues within the same protein pair and residues from random protein pairs show mean RMSDs of 21.7 Å and 23.2 Å, respectively, as can be seen in Figure 3C. The model actively differentiates structurally conserved regions from dissimilar ones, and the representational signal tracks this geometric discrimination faithfully.

This performance likely reflects a combination of MSA support and, given that all SABmark proteins predate the AlphaFold 3 training cutoff, possible prior exposure to these folds. To disentangle these contributions and directly assess the role of co-evolutionary information, we reran the identical analysis without MSA input. The contrast is immediate: experimentally aligned regions now yield a mean RMSD of 12.8 Å, no longer showing meaningful structural concordance, while control RMSDs increase only modestly to 23.2 Å and 24.5 Å, as shown in Figure 3C.

Importantly, as shown in Figures 3D and 3F, this loss of structure in the no-MSA setting does not improve the structural alignment of metamorphic sites with their experimental structures. Furthermore, Figure 3A shows that the representational gap between true alignments and within-pair random controls narrows substantially, indicating a marked reduction in the model’s ability to distinguish genuinely conserved structural regions from irrelevant residue matches.

This structural collapse is mirrored in the latent space. The distinct representational clustering that separated structurally related groups under the MSA condition dissolves in the no-MSA setting, the effective rank (see Methods) of the C_N_ pair representations, for example, rises from about 15.2 with MSAs to 24.2 without them, indicating a less organized latent space with weaker geometric separation between fold groups (Figure 3A). AlphaFold 3 can identify structurally overlapping regions with remarkable precision among sequence-dissimilar proteins—but only when MSA information is present. The failure to recover even canonical fold overlaps without an MSA, regardless of potential training familiarity, establishes co-evolutionary information as a dominant substrate of the model’s structural reasoning.

### 2.8 AlphaFold 3 Collapses Metamorphic Sites to a Single Fold

To test the opposite regime from SABmark, we asked how AlphaFold 3 behaves when sequences are highly similar but experimentally validated structures differ. We assembled 46 protein pairs from a previous fold-switching study [23], each containing experimentally validated regions known to adopt alternative folds—a stringent test of whether AlphaFold 3 can distinguish sequence-similar regions that should map onto distinct structural states.

Structures were predicted and compared specifically over the regions known to adopt different folds. As in the SABmark analysis, two controls were used: random regions of equivalent length from the same protein pair as an internal control, and random regions from random monomeric proteins in the 400-protein benchmark as an external null (see Methods).

Under the MSA-conditioned setting, the result is unambiguous. The fold-switching regions are predicted to be far more structurally similar to one another than they should be experimentally, with a mean RMSD of only 2.02 Å across aligned regions, compared with 9.25 Å and 10.3 Å for the within-pair and random-monomer controls, respectively. The representational analysis reinforces this conclusion: cosine similarity between aligned fold-switching regions is substantially higher than in either control condition, indicating that the latent space strongly favors a shared fold hypothesis across regions that experimentally should diverge. This conformational bias is not a property of the final diffusion module alone—it is already embedded in the pair representations at C_1_ and C_N_, consistent with the Pairformer’s tendency to converge toward a single dominant geometric hypothesis.

An important nuance emerges from the uncertainty profile. Although AlphaFold 3 converges toward the same dominant fold, it partially recognizes the fold-switching regions as anomalous. Mean distogram entropy in fold-switching regions is 2.25, compared with 1.39 in non-switching regions of the same proteins, mirrored by a lower mean predicted Local Distance Difference Test (pLDDT), which is an AlphaFold 3 confidence metric that predicts atomic orientation [4], of 77.4 versus 82.3 elsewhere. The model is aware that something is geometrically unusual at these sites, even while failing to resolve the alternative conformation.

The mechanistic role of the MSA becomes clear when it is removed. Without MSA input, fold-switching regions exhibit substantially higher pairwise RMSD relative to one another, but this does not reflect improved resolution of alternative conformations. It reflects generalized structural inaccuracy. MSA-conditioned predictions of the fold-switching regions have a mean RMSD of 7.35 Å relative to the experimental structure; no-MSA predictions degrade further to 10.0 Å. Removing the MSA does not make the model more conformationally flexible—it makes it less correct (see Figures 3D and Figures 3F).

This places the fold-switching benchmark within the same pattern established across the 400-protein benchmark and SABmark. AlphaFold 3 recovers a stable and often accurate dominant fold even at metamorphic sites when MSA information is present. Without it, performance collapses regardless of whether the challenge is sequence novelty, mutational load, sequence dissimilarity, or experimentally validated conformational switching. That this degradation occurs even for proteins likely familiar from training indicates that the model’s structural prior is organized around MSA-derived evolutionary context, not raw sequence—a conclusion we now test directly through systematic perturbation of the MSA itself.

### 2.9 Structural Accuracy Requires MSA Presence, Not Depth or Integrity

Having observed persistent structural and representational degradation when MSAs are absent, we next asked how the quality and quantity of MSA information shape model performance across the continuum between full presence and complete removal. To systematically probe this continuum, we performed two complementary perturbation analyses on the 200 novel proteins. In the first, an increasing fraction of MSA columns was randomly shuffled at levels of 10, 20, 40, 70, 90, 95, and 99%, testing sensitivity to corruption of co-evolutionary signal at fixed depth. In the second, MSAs were subsampled at the same fractional levels, independently measuring sensitivity to loss of alignment depth while preserving co-evolutionary integrity (see Methods).

The results are remarkably robust. As shown in Figures 4A–F, neither subsampling nor column shuffling substantially decreases structural accuracy, representational stability, distogram entropy, or the effective rank of the pair representations until perturbations become extreme. Structural performance remains largely intact until more than 99% of the MSA has been removed—leaving on average only 47 sequences—and until more than 40% of MSA columns have been shuffled. Even under the most severe perturbations tested, mean global fold quality remains well above 0.5, indicating that predicted structures still preserve accurate overall topology. As shown in Figures 4C and 4D, the cosine distance from the original prediction is high at C_1_, reflecting substantial representational divergence at early checkpoints, but drops sharply by C_N_, indicating that the Pairformer recovers a structurally coherent fold even from the diminished signal. Taken together, these perturbation results reinforce the near-binary character of MSA dependence identified in the preceding analyses: what matters for model performance is not the quantity or integrity of evolutionary information but simply its presence.

**Fig. 4.**
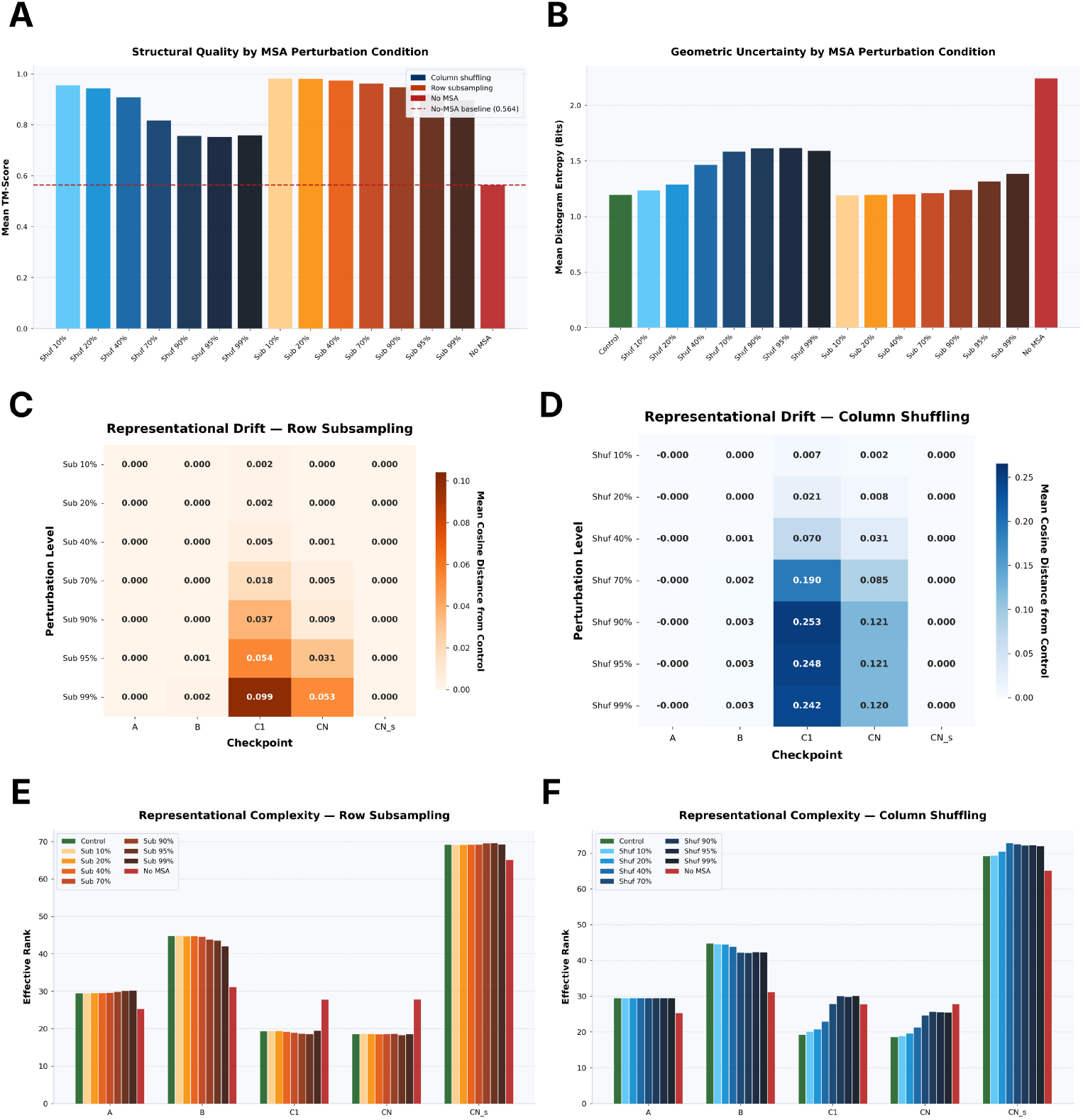
Structural quality and representational geometry under MSA perturbation. **(A)** Mean TM-score across all column-shuffling and row-subsampling perturbation levels relative to the no-MSA baseline, demonstrating that even sparse or informationally degraded MSAs substantially preserve structural prediction quality. **(B)** Mean distogram entropy across the same perturbation levels relative to the noMSA baseline, showing that geometric uncertainty remains markedly lower than the no-MSA condition even under aggressive perturbation. **(C–D)** Heatmaps of mean cosine distance from the unperturbed control across all pair and single checkpoints (*A, B, C*_1_, *C*_*N*_, *C*_*N*_ -single) for row subsampling (C) and column shuffling (D). Both perturbation types induce relatively modest representational drift, indicating that the internal representations are largely invariant to MSA content degradation of this kind. **(E–F)** Entropy-based effective rank of the representational geometry across all perturbation levels for row subsampling (E) and column shuffling (F). The effective rank is substantially elevated in the no-MSA condition for the later pair checkpoints, indicating that the absence of MSA produces a more distributed and less consolidated representational space.

### 2.10 Phylogenetic Subsampling Reveals the Necessity for MSA Sequence Diversity

A natural question arising from this is whether the content of the MSA matters— and if so, what properties of evolutionary context are most informative. To investigate this, we performed phylogenetic subsampling of the 200-protein dataset, constructing twelve variants of each protein’s MSA restricted to depths of 1, 5, and 10 sequences drawn from one of four phylogenetic tiers: sequences with greater than 80% identity to the query (similar), sequences with 50–70% identity (medium), sequences with less than 30% identity (dissimilar), and sequences drawn uniformly at random. Due to the identity distribution constraints imposed by real MSAs, only 61 of the 200 proteins had sufficient coverage across all three non-random tiers to qualify for analysis, and all subsequent results pertain to this subset.

The results reveal that MSA content matters profoundly, and in ways that are not predicted by classical models of co-evolutionary inference. First, structural accuracy and model confidence rose markedly from 1 to 10 sequences across all tiers, demonstrating that even a single homologous sequence provides the model with sufficient evolutionary context to dramatically improve over the no-MSA baseline (Figures 5A and 5B). Given the binary-threshold behavior identified above, this is consistent with a model that treats evolutionary context categorically, where the presence of a homologous sequence activates structural priors encoded in the model weights, while its absence leaves the model without the reference frame it requires.

**Fig. 5.**
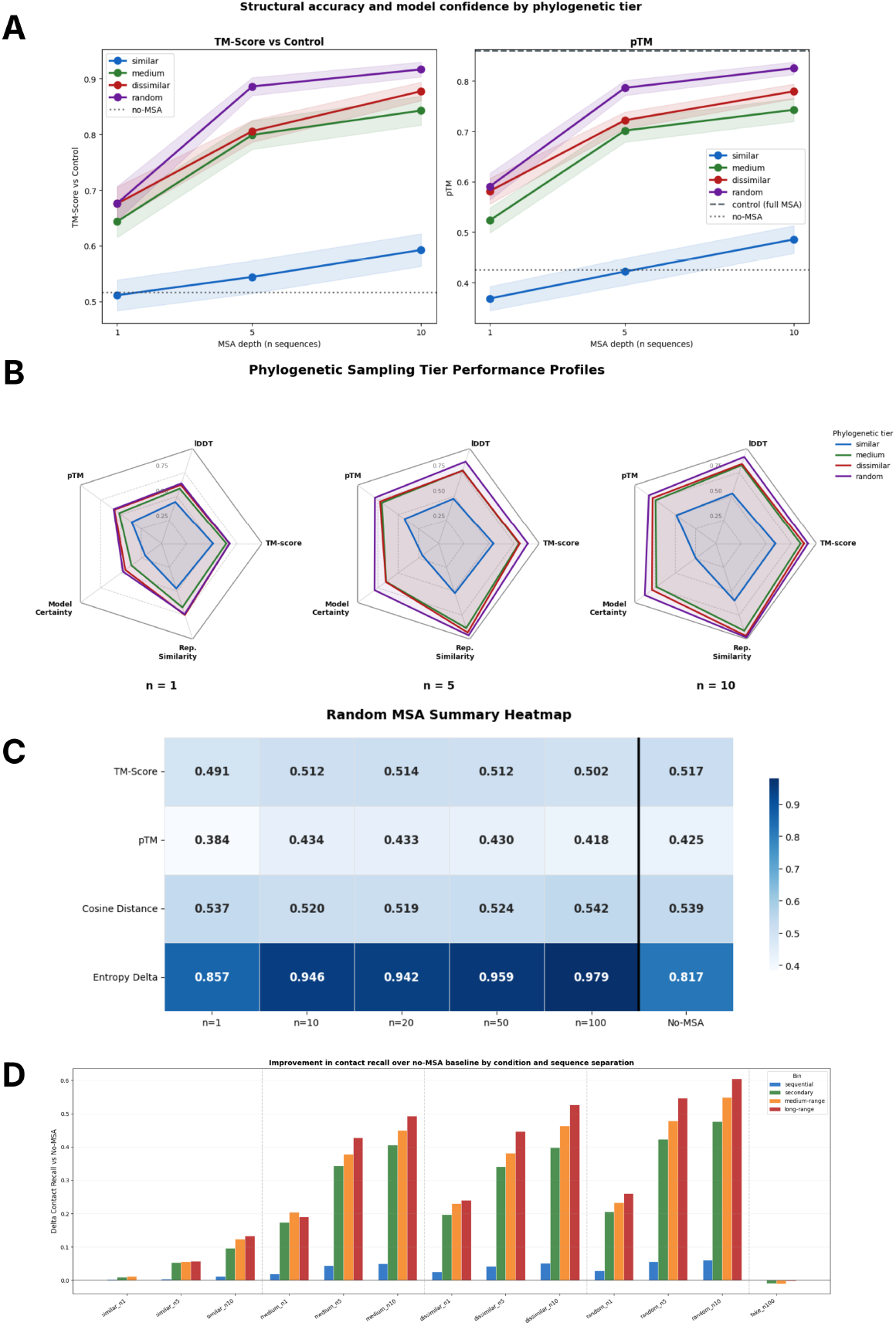
Phylogenetic diversity of MSA sequences determines structural accuracy and representational fidelity in AlphaFold 3. **(A)** Line plots showing the improvement in TM-score and pTM relative to the full-MSA control prediction as a function of MSA depth, stratified by phylogenetic tier. Greater sequence diversity yields greater improvements across all depths, with randomly selected sequences outperforming all curated tiers at matched depth. **(B)** Performance profiles illustrating model accuracy and internal representational quality across five axes—TM-score, lDDT [37], pTM, 1 - Entropy Increase (model certainty), and 1 - *C*_*N*_ Cosine Distance (representational similarity to the full-MSA prediction)—for each phylogenetic tier at MSA depths of 1, 5, and 10 sequences. A larger enclosed area indicates superior performance across all metrics simultaneously. Sequences with low homology to the query consistently produce broader profiles than near-identical sequences, particularly as depth increases. **(C)** Summary heatmap of model performance under injection of fake MSA sequences constructed from randomly truncated alignments of unrelated proteins. Across all metrics and all depths tested, fake sequences fail to improve over the no-MSA baseline and produce elevated distogram entropy. **(D)** Delta *C*_*α*_–*C*_*α*_ contact recall relative to the no-MSA baseline, binned by sequence separation. Labels denote phylogenetic tier and MSA depth: *similar n#* indicates sequences with ≥80% identity to the query, *medium n#* sequences with 50–70% identity, *dissimilar n#* sequences with ≤30% identity, *random n#* uniformly sampled sequences, and *fake n100* the unrelated-protein MSA control. Long-range contacts (≥ 24 residues apart) show the largest and most consistent improvements with increasing phylogenetic diversity, reflecting the dependence of tertiary structure on co-evolutionary information that near-identical sequences cannot provide.

Second, and more remarkably, sequences highly similar to the query—despite being the closest evolutionary relatives—provided the least benefit of any tier, with structural accuracy and model confidence remaining near no-MSA levels even at *n* = 10 (Figures 5A and 5B). This is a counterintuitive result: near-identical sequences, which are considered the most informative homologs in classical comparative genomics, actively fail to improve AlphaFold 3’s performance. We interpret this as evidence that the model requires phylogenetic diversity rather than phylogenetic proximity. Near-identical sequences are informationally redundant—they confirm what the query sequence already encodes and provide no differential signal about which positions are structurally constrained and which are variable. Diverged sequences carry the evolutionary record of what can and cannot change, which appears to be precisely the signal the model needs.

Third, randomly selected sequences—which sample the identity spectrum naturally, with a mean aligned similarity of approximately 0.42 and standard deviation of 0.14 across all depth thresholds—outperformed all phylogenetically constrained tiers at matched depth, most strikingly at *n* = 10 where random selection produced accuracy and confidence approaching the full-MSA control. This is not because random sequences are intrinsically superior, but because they incidentally capture the diversity that the model requires: by sampling without regard to identity, random selection spans the full phylogenetic spectrum and provides the breadth of evolutionary signal that any single tier constrains away.

Fourth, to determine whether the MSA must carry any biophysical relevance to the query sequence at all, we injected sequences drawn from the MSAs of the 139 non-qualifying proteins into each of the 61 qualifying proteins’ input features, at depths of 1, 10, 20, 50, and 100 sequences, trimming each donor sequence to match the target alignment length (see Methods). As shown in Figure 5C, the performance of these fake sequences is no better than the no-MSA baseline across all metrics— structural accuracy, model confidence, cosine distance, and distogram entropy—and does not improve with increasing depth. This null result is a critical complement to the phylogenetic subsampling findings: MSA existence alone is insufficient, and the signal AlphaFold 3 extracts from the MSA is genuinely evolutionary. The model is not reading alignment-shaped input.

To probe which structural features drive the TM-score improvements observed across phylogenetic conditions, we computed pairwise *C*_*α*_–*C*_*α*_ contact maps for all 61 proteins under each condition and compared them against the full-MSA control prediction. Contacts were assigned to four bins by sequence separation: sequential (1– 5 residues), secondary (6–11), medium-range (12–23), and long-range (≥ 24). For each protein and bin, we report the delta contact recall relative to the no-MSA baseline, isolating the improvement attributable to MSA inclusion from any protein-intrinsic baseline recovery.

As shown in Figure 5D, the inclusion of even minimally diverse MSA sequences produces marked improvements in contact recall across all separation ranges, with long-range contacts consistently showing the largest gains. Long-range contacts are precisely those that cannot be inferred from local sequence patterns alone and require comparative evolutionary information. The model’s ability to recover them with as few as one dissimilar or randomly selected sequence supports the view that AlphaFold 3 has internalized co-evolutionary constraints during training and requires only a minimal evolutionary signal to activate them at inference time. Random sequences produce the largest contact recall gains at all separation ranges and depths, while similar sequences yield improvements that are marginal even at *n* = 10, reinforcing that proximity to the query sequence is not a proxy for informational content. The fake MSA condition produces negligible or negative delta recall, confirming that evolutionary relevance— not alignment format or depth—is what drives contact recovery.

### 2.11 Structural Accuracy Requires Phylogenetic Diversity, Not Alignment Depth

To understand the mechanistic basis of this diversity requirement at residue resolution, we performed a residue-level analysis of long-range contact recovery at *n* = 1. Long-range contacts are critical for stabilizing protein structure and enabling correct folding [38, 39], and our previous analyses showed that they benefit most from increased MSA diversity and depth. This setting is analytically tractable in a way that larger depths are not. At *n* = 5 and *n* = 10, the combinatorial space of residue-pair interactions across multiple homologs obscures the contribution of individual comparisons, while the sequence count remains too small for reliable covariation statistics.

At *n* = 1, the alignment reduces to a direct binary comparison between the query and a single homolog: each residue is either conserved (identical to the query) or variable (differing from the query or gapped), yielding three classes of long-range pairwise contacts—conserved–conserved (CC), conserved–variable (CV), and variable– variable (VV).

The residue composition of the *n* = 1 alignment differs markedly across tiers and is central to interpreting the results. The similar tier contains on average 27.6% variable residues, whereas the dissimilar and random tiers contain 86.2% and 73.3%, respectively. In the similar tier, conserved residues largely reflect phylogenetic inertia: insufficient evolutionary time has elapsed to test which positions are truly essential. By contrast, a residue that remains conserved across 70% sequence divergence has survived sustained purifying selection and is a reliable indicator of structural necessity.

These differences are directly reflected in contact recovery. Long-range CC recall is 0.315 in the similar tier, rising to 0.675 in the dissimilar tier and 0.671 in the random tier. This improvement does not arise from a greater abundance of CC contacts— CC pairs account for only 3.6% of long-range control contacts in the dissimilar tier compared to 65.4% in the similar tier. Instead, conserved residues in distant homologs are structurally informative in a way those in near-identical sequences are not. They mark the positions that define global topology, and their long-range contacts provide the scaffold upon which the rest of the fold is organized.

The ordering of recall classes within the diverse tiers confirms this scaffolding mechanism. In the dissimilar tier, CC recall (0.675) exceeds CV recall (0.601), which exceeds VV recall (0.514). The same ordering holds in the random tier: CC 0.671, CV 0.627, VV 0.454. The model resolves contacts between genuinely conserved residues most reliably. These anchor interactions establish the geometric context within which contacts involving variable positions become interpretable. Variable residues—despite comprising the large majority of the alignment in diverse conditions—do not drive contact recovery autonomously.

The similar tier fails at every level of this hierarchy. With 65.4% of long-range contacts falling between residues conserved only through phylogenetic inertia, the model receives no differential signal about which positions are structurally critical. CC recall remains at 0.315—near the no-MSA baseline—because conservation in a near-identical sequence carries no information about structural necessity. Without reliable anchors, neither conserved nor variable contacts can be confidently resolved. This failure is not a matter of insufficient depth or alignment format: it reflects a fundamental absence of the evolutionary contrast needed to distinguish structurally essential from permissive positions.

Together, these findings point toward an important practical principle: structural accuracy does not require deep alignments, but it does require sequences that have diverged sufficiently to expose which positions are truly constrained. To test whether this principle generalizes beyond the 61 qualifying proteins, we implemented a nested subsampling protocol restricted to sequences with 30–60% identity to the query— the range that corresponds to the natural identity distribution of randomly selected sequences, which performed best across all preceding analyses. Of the 200 novel proteins, 183 had MSAs sufficiently deep to support this analysis (at least 10 sequences in the target identity range). Relative to full-MSA control predictions, these shallow mid-homology MSAs achieved a mean TM-score of 0.88 (median 0.94) at *n* = 5, rising to a mean of 0.92 (median 0.97) at *n* = 10, demonstrating that a handful of appropriately diverged sequences is sufficient to recover the large majority of structural accuracy achievable with a full evolutionary alignment.

## 3 Discussion

This work represents, to our knowledge, the first systematic mechanistic interpretability study of AlphaFold 3’s internal representations. Prior examinations of AlphaFold-like systems have largely remained at the phenomenological level, characterizing outputs without probing the internal geometry that produces them [10, 13, 21, 23, 40, 41]. By extracting and analyzing the single and pair representations across four checkpoints, we trace the progressive reorganization of the latent space from a sequence-length proxy at Checkpoint A to a linearly decodable, geometrically coherent representation of three-dimensional structure at Checkpoint C_N_. The pair representations are particularly rich, encoding residue contacts, pairwise distances, secondary structure, and global confidence in a form that becomes increasingly accessible as the Pairformer and recycling steps refine the embedding manifold. That this refinement follows a consistent, compressive trajectory into a lower-dimensional effective feature space suggests that substantial information remains to be extracted from these internal states.

The activation patching experiments open a compelling direction. By showing that the dominant axes of the pair representation causally govern downstream distogram confidence—and that representations encoding successful folds can be transplanted across unrelated proteins—we demonstrate that AlphaFold 3’s latent geometry is not merely descriptive but manipulable. This raises a question the framework makes tractable but that this study does not resolve: whether direct intervention on internal representations can induce structures the model would otherwise suppress due to evolutionary precedent or data scarcity. These representations may also provide richer uncertainty quantification than scalar metrics such as pLDDT or pTM, as their high-dimensional geometry encodes early signals of representational failure before it manifests in predicted coordinates.

This representational framework also reveals the central and perhaps underappreciated role of the MSA in a way that output-level analyses cannot. While it is broadly understood that shallow MSAs degrade performance [4, 40, 41], the internal representation analysis developed here reveals that MSA absence does not merely reduce structural accuracy: it destroys internal representational coherence entirely. AlphaFold 3’s ability to identify structurally similar regions across sequence-dissimilar proteins is nullified without the MSA. Its mutational invariance—both structural and representational—depends on MSA availability. For fold-switching proteins, the model is conformationally biased with an MSA, but without one it cannot even recognize conformational similarity informed by sequence. This pattern holds across every benchmark tested, including proteins likely within or similar to the AlphaFold 3 training set. Without an MSA, AlphaFold 3 fails even for small, simple monomers, and the internal representations collapse with it. For multimeric complexes, which are far more structurally challenging and whose interactions govern much of biology, the situation is unlikely to improve [21, 26, 42]. Sequence alone, regardless of similarity to training examples, is not sufficient. The model appears to have learned a mapping from MSA-derived motifs—perhaps coupled with sequence—to structure, and that mapping dominates.

From a purely epistemic standpoint, this is cause for reflection. AlphaFold 3 is immensely powerful, but its power is bounded by our ability to construct informative MSAs. Whatever physical understanding the model has acquired is overwhelmingly mediated by evolutionary statistics and their encoded echo rather than autonomous biophysical reasoning. This is a working solution to the protein structure prediction problem, but it does not reflect how proteins actually fold. When a protein folds in nature, it does not query an evolutionary database. It follows biophysical principles and minimizes free energy. The folding process is governed by the physics of the polypeptide chain in its environment, not by the statistical preferences of its homologs. A model that recovers structure primarily through evolutionary context and the implied similar fold is solving a related but distinct problem, and solving it brilliantly is not the same as understanding it.

However, this dependence is less restrictive than it initially appears. The MSA perturbation experiments show that AlphaFold 3 tolerates substantial corruption and depth reduction, maintaining structural accuracy under column shuffling and subsampling. The model does not require deep or pristine alignments—only sufficient diversity to anchor the Pairformer’s geometric reasoning. Indeed, as few as 5–10 evolutionarily divergent sequences relative to the query may suffice.

This explains why even highly nonphysical mutations fail to disrupt the predicted fold. When up to 40% of residues are replaced without regenerating the MSA, the alignment still encodes the original evolutionary motifs, constraining the model to the same structural basin. Critically, even when MSAs are regenerated from these mutated sequences, the predicted structure shows no substantive deviation from the original fold [21]. Although the query is corrupted, the regenerated MSAs still contain enough diverse, distant homologs to meet the model’s minimal diversity requirement. Thus, whether the MSA is fixed or regenerated, the same fold is recovered: in the former case due to preserved coevolutionary signal, and in the latter due to sufficient residual diversity. The lack of any meaningful difference between these regimes reinforces that AlphaFold 3 functions less as a precise sequence-to-structure mapper and more as a sophisticated fold recognition system driven by a small set of phylogenetically diverse sequences.

This observation suggests a practical path forward. Rather than relying on expensive MSA collation, one can instead identify minimal, optimally diverse sequence sets. This directly supports emerging synthetic MSA generation methods [43], which aim to construct informative alignments for proteins with sparse natural homologs. If heavily degraded natural MSAs preserve predictive accuracy, well-designed synthetic ones should as well—provided they capture sufficient diversity. The representational framework developed here offers a principled way to evaluate whether any MSA, synthetic or natural, maintains the coherent internal geometry required for accurate prediction.

Broadly, these results motivate methods that leverage minimal evolutionary scaffolds to achieve accurate, generalizable prediction, which would extend structural modeling into protein space that evolutionary history has not yet sampled [44, 45].

To truly harness structural prediction for scientific progress, especially as we seek to design new proteins or explore regions of protein space beyond the reach of evolutionary sampling, we cannot remain bounded by the richness of evolutionary history. The protein structure prediction problem, as historically defined, has been largely solved [46]. The deeper problem—a physically grounded, genuinely generalizable model of how sequence determines structure—remains open. The work presented here does not close it, but it sharpens the terms of the problem: a model that genuinely reasons from sequence to structure would not need evolutionary scaffolding to reach that geometry—it would reach it from first principles. The internal representations developed here offer a principled framework for measuring progress toward that goal, and for detecting, concretely, when it has been achieved.

## 4 Methods

### 4.1 Dataset Collection

The primary dataset used in this study is composed of 400 monomer proteins—200 of which are called ‘Similar’ and 200 of which are called ‘Novel’ by virtue of their sequence similarity to proteins in the PDB released on or before September 30, 2021 [5, 47, 48]. Proteins labeled as Similar share 30% or more sequence homology with pre-September 2021 PDB entries, meaning their homologs could plausibly have been in AlphaFold 3’s training set [21]. In contrast, Novel proteins have no such homology. Importantly, all proteins in the dataset were released after September 30, 2021, and no two proteins share 30% or more sequence similarity, ensuring minimal redundancy and that none were present in AlphaFold 3’s training set [5, 47]. This categorization enables a detailed assessment of AlphaFold’s performance across different protein types and levels of novelty.

Two additional datasets were used in this study: SABmark and a dataset of experimentally validated fold-switching proteins from [23, 49]. The SABmark subset comprises 100 proteins organized into 5 groups of 20. Within each group, proteins belong to the same Structural Classification of Proteins (SCOP) fold category [50], ensuring high experimentally validated structural similarity, while maintaining low sequence homology (≤ 25% pairwise identity) [49]. This design enables the analysis of whether internal representations preserve fold-level geometric organization despite minimal sequence similarity. Across the 5 groups, the full set contains 950 possible within-group protein pairs. Of these, 560 pairs include complete residue-level structural overlap annotations and were retained for downstream analysis—any pairs lacking full pairwise alignment annotations or other information were excluded. To ensure architectural diversity, each of the five groups was selected from a distinct SCOP group, with no PDB structure appearing in more than one group [50].

The fold-switchers dataset comprises 46 pairs filtered to isolate high-quality metamorphic transitions [23]. To prevent massive static domains from masking the metamorphic signal, pairs were excluded if the total chain length exceeded ten times the fold-switching region. We also removed redundant PDB structures and required at least 60% local sequence identity between the annotated switching fragment and the deposited PDB sequence. This pipeline yielded 46 robust pairs, each capturing two distinct, experimentally validated conformational states for the same underlying sequence.

### 4.2 Embedding, Structure Extraction and Standardization

We extract single (**s** ∈ ℝ^*L*×384^) and pair (**p** ∈ ℝ^*L*×*L*×128^) representations at four checkpoints spanning the AlphaFold 3 forward pass [4]. Checkpoint A captures sequence initialization with shallow co-evolutionary statistics from the MSA (per-position amino acid frequencies and deletion means). Checkpoint B is sampled immediately after the MSA module, introducing deeper evolutionary context via outer product mean and row-wise self-attention. Checkpoint C_1_ follows the first Pairformer pass–through, representing the earliest point of structural self-organization. Checkpoint C_*N*_ follows all Pairformer layers and recycling iterations, representing the mature representation before the diffusion model.

For analyses comparing representations across proteins of different lengths, we standardize the embeddings via global mean pooling to yield fixed-dimensional vectors per protein [11, 51]. For the AlphaFold 3 single representation, we compute the mean across the sequence dimension:

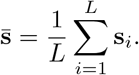

For the pair representation, we apply global upper-triangular mean pooling. By extracting all unique residue pairs (*i, j*) where *i < j* and *i, j* are residue indices, we compute the mean across the spatial dimensions:

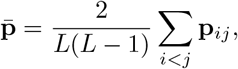

yielding a single 128-dimensional vector. While pooling inevitably discards fine grained pairwise geometric detail, it is necessary for length-invariant comparison across diverse datasets; other alternatives, such as padding, truncation, or length-matching, introduce greater structural artifacts [11, 51, 52]. For analyses on identical proteins under perturbation—SABmark fold clustering, mutational studies, fold-switching—we preserve the full unpooled representations with no information loss.

We also evaluated alternative feature aggregation and augmentation strategies, such as mean+std pooling for global representations and a context-augmented approach that incorporates local residue-level statistics and sequence separation for pairwise features. However, as illustrated by the representative comparison between raw pairwise features and the context-augmented strategy for *C*_*α*_ − *C*_*α*_ distance prediction in Figure 1L, these more complex schemes did not majorly or consistently outperform simpler representations.

Critically, across all generated structures and the embeddings that precede them, no structural templates were used. As is mentioned in further detail, some analyses included the MSA and some did not as inputs into AlphaFold 3, but no analysis used template information as inputs to reduce potential confounds and structural invariance.

For the purposes of structure generation, all default settings of AlphaFold 3 were employed with one exception. Instead of generating five diffusion samples, as is standard, for computational efficiency only one diffusion sample was generated during structural inference.

### 4.3 Linear Probing and Structural Feature Analysis

#### 4.3.1 Linear Ridge Regression Probes

To test whether AlphaFold 3 representations encode specific structural or biophysical properties, we train linear probes on per-residue or per-pair feature vectors. For regression tasks, which operate on continuous structural properties, we use Ridge regression with *ℓ*_2_ regularization [30]. For a given representation (single or pair), we standardize the feature matrix via:

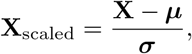

where **X** is the feature matrix, ***µ*** is the vector of column-wise feature means, and ***σ*** is the vector of column-wise standard deviations, and fit a Ridge model with regularization parameter *α* = 1.0. Test performance is measured via the coefficient of determination *R*^2^:

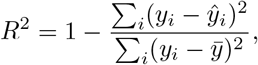

which quantifies the proportion of variance in the target variable explained by the probe, with values closer to 1 indicating stronger predictive performance and values near 0 indicating performance no better than predicting the mean

Across the board, we use a consistent 80/20 train/test split on the 400-constituent protein dataset. To strictly prevent data leakage, this split is performed at the whole protein level. To ensure both sets accurately represent the dataset’s distribution of structural quality, the selection was stratified by predicted Template Modeling (pTM) score, where proteins were first categorized into discrete bins based on their pTM, and the random 80/20 split was applied independently within each bin to prevent evaluation bias from a skewed test set. Furthermore, because pairwise representations scale quadratically with sequence length (*O*(*L*^2^)) and quickly exceed practical memory limits during model fitting when utilizing raw *i, j* pair feature vectors, we uniformly subsampled a maximum of 3,000 residue pairs per protein using a deterministic seed for exact reproducibility. Prior to fitting, standardization was performed using the `StandardScaler` function from the *Scikit-Learn* package [53]. Model performance is evaluated using the coefficient of determination (*R*^2^).

In addition to Ridge regression, we tested Multilayer Perceptrons (MLPs) and Gradient Boosting Trees (GBTs). All methods performed comparably, and, ultimately, the Ridge Regression method was chosen due to its simplicity, interpretability, and regularization penalties that minimize overfitting [54–56].

#### 4.3.2 Biophysical Feature Extraction from PDB Structures

Biophysical labels were derived directly from PDB coordinate files using standard structural analysis pipelines. For each residue, we computed complementary structural properties spanning local conformation, solvent exposure, and pairwise geometry. Secondary structure assignments were obtained using a hierarchical fallback strategy. We first extracted annotations from mmCIF_`struct_conf` and _`struct_sheet_range` records, which provide helix and sheet labels. When these were unavailable, we applied *DSSP* to the *C*_*α*_ trace to classify residues as helix, strand, or coil [57]. Per-residue solvent-accessible surface area (SASA) was computed using the Shrake–Rupley algorithm with a probe radius of 1.4 Å, yielding values in Å^2^ [58]. To complement absolute solvent exposure, we defined a normalized burial score based on local neighborhood density. For each residue *i*, we counted the number of residues *c*_*i*_ whose *C*_*α*_ atoms lie within 10 Å:

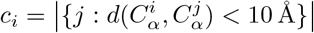

We then identified the *k* = 9 nearest neighbors of residue *i* in *C*_*α*_ distance space, denoted as NN_9_(*i*), and summed their counts:

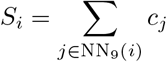

The final burial score was defined as:

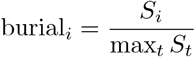

yielding values in [0, 1]. Unlike SASA, this measure emphasizes relative local packing density and is less sensitive to absolute solvent-exposure scale. For pairwise structural properties, we computed full *C*_*α*_–*C*_*α*_ distance matrices:

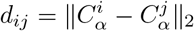

Contact maps were then derived as binary indicators:

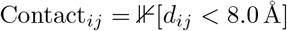

Biophysical labels were available only for structurally resolved residues, meaning that analyses involving PDB-derived features were restricted to this subset of residues in each protein.

#### 4.3.3 Principal Component Analysis and Effective Dimensionality

To assess the intrinsic, structured dimensionality of representations across checkpoints, we fit `IncrementalPCA` on standardized training data [53]. We compute the explained variance ratio *λ*_*k*_ for each principal component *k*, then determine the effective dimensionality as the number of components required to explain 95% of total variance: 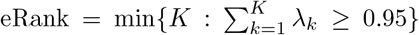 [32]. Lower effective dimensionality indicates more compressed, structured geometry; higher rank indicates distributed information. We also compute Spearman rank correlations between individual principal component scores and structural labels to identify which components encode specific biophysical features. By projecting pooled representations into this principal subspace, we can track how the model’s internal representations reorient from sequence-related features toward biophysical and structural descriptors as it progresses through the Pairformer layers.

#### 4.3.4 Unique Information Decomposition

To understand the division of labor between architectural tracks, we decompose the information content of the single and pair representations into unique and shared components. Unlike the other linear probes, this analysis focuses on residue-level properties. We therefore mean-pool the pair representation matrix along one spatial dimension to convert the *O*(*L*^2^) pair data into an *O*(*L*) residue-level neighborhood representation. This reduction in computational scaling allows the analysis to utilize all of the residues in the dataset, bypassing the memory constraints that necessitate subsampling in pair-to-pair tasks.

For a given structural property and checkpoint, we train three Ridge regression probes:

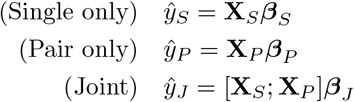

where **X**_*S*_ ∈ ℝ^*L*×384^ is the per-residue single representation, **X**_*P*_ ∈ ℝ^*L*×128^ is the pooled pair representation, and [**X**_*S*_; **X**_*P*_ ] denotes their concatenation. Let 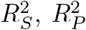, and 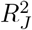 denote the test *R*^2^ for each model, calculated using the same stratified 80/20 protein-level split as the previous experiments to ensure generalization to unseen structures. We then define the information decomposition as:

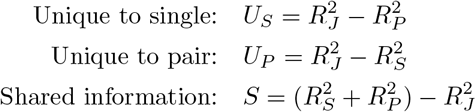

A positive *U*_*S*_ indicates that the single representation encodes structural information not present in the pair track, while *U*_*P*_ quantifies the unique contribution of the pair representations. Positive *S* values denote redundancy, indicating that both representations capture overlapping views of the same structural property.

### 4.4 Distogram Entropy and KL Divergence Analysis

The distogram—the predicted pairwise distance distribution—encodes geometric uncertainty at each residue pair [3, 4, 47]. We extract the softmax-normalized distance predictions **p**_*ij*_ ∈ ℝ^*C*^ (where *C* is the number of distance bins, which is 64, as is the default) for all pairs (*i, j*) at each checkpoint. We compute the Shannon entropy of each pairwise distribution:

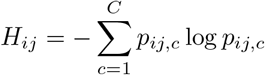

Higher entropy indicates more distributed predictions, which shows greater uncertainty. Lower entropy indicates denser and more certain predictions. We stratify analysis by sequence separation, *s* = |*j* −*i*|, defining four bins: sequential (1–5 residues apart), secondary (6–11), medium-range (12–23), and long-range (≥ 24). For each separation bin and each protein, we compute the mean entropy in the MSA and no-MSA conditions, then the entropy difference Δ*H* = *H*_no-MSA_ − *H*_MSA_. A positive Δ*H* indicates that the MSA reduces uncertainty, meaning it makes distance predictions more geometrically certain.

To measure the magnitude of representational change induced by the MSA, we compute the Kullback–Leibler (KL) divergence from MSA to no-MSA distributions:

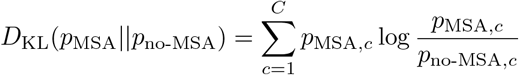

We compute this per-pair and then average within each sequence-separation bin, testing whether the MSA’s influence on pairwise geometry is strongest at long-range contacts (which are typically underdetermined by sequence context alone) or uniform across all separations.

### 4.5 Effective Rank

In later analyses, we also employ an alternative measure of intrinsic dimensionality: the effective rank, defined as the exponential of the Shannon entropy of the normalized singular value spectrum,

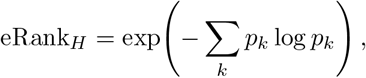

where *p*_*k*_ = *σ*_*k*_*/* ∑_*j*_ *σ*_*j*_ are the singular values normalized to sum to one [36, 59]. Unlike the variance-threshold measure, this quantity integrates continuously over the full singular value spectrum and is sensitive to how concentrated or diffuse the variance is across all dimensions simultaneously, rather than whether a fixed cumulative threshold is crossed. The two measures are qualitatively consistent across the checkpoints and conditions analyzed here, and where each is used is indicated explicitly in the text.

#### 4.5.1 Distance Decodability Probes

To assess how linearly pair representations encode pairwise residue distances, we trained three Ridge regression probes predicting ground-truth *C*_*α*_–*C*_*α*_ distance *d*_*ij*_: a sequence-separation baseline using only |*i* −*j*|; an isolated pair probe using **p**_*ij*_ directly; and a context-augmented probe that additionally appends the row and column means of the pair representation matrix alongside |*i* − *j*|, incorporating each residue’s global structural neighborhood. All probes were evaluated using the 80/20 train-test split used in prior probes and 3000 residue pairs were uniformly subsampled.

### 4.6 Activation Patching and Latent Confidence Modulation

To establish a causal link between the Pairformer latent space and the network’s structural confidence, we employ activation patching on the C_N_ checkpoint pair representation **P** ∈ ℝ^*L*×*L*×128^ immediately preceding the distogram head. Unlike other structural modules, the AlphaFold 3 distogram head acts as a direct, transparent linear probe [4]. For a given residue pair (*i, j*) with representation vector **p**_*ij*_ ∈ ℝ^128^, the pre-softmax logits **z**_*ij*_ ∈ ℝ^64^ are computed via a learned weight matrix **W** and bias **b**, followed by symmetrization [4]:

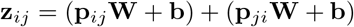

Because this projection lacks intermediary non-linearities or layer normalization, targeted interventions on **p**_*ij*_ map directly to changes in the output probability distribution and its corresponding Shannon entropy *H*_*ij*_, allowing us to isolate axes of geometric confidence.

#### 4.6.1 Identifying Axes of Variance

To identify the principal directions of variation within the latent space, we applied `IncrementalPCA` to the unpooled, upper-triangular pair representations. Crucially, the dataset was first split into training and testing sets, stratified across ten TM-score deciles. This stratification guarantees that the PCA was fit on an equal representation of highly confident (accurate) and highly uncertain (failed) structural geometries. This procedure yielded principal direction vectors **v**_*k*_ and their corresponding eigenvalues *λ*_*k*_.

#### 4.6.2 Modulating Geometric Confidence

To determine if these principal components act as causal “dials” for structural certainty, we mathematically modulated the representations of target proteins drawn strictly from the held-out test set (comprising the 40 highest and 40 lowest TM-score predictions). For a chosen principal component *k*, we shifted the target protein’s pair representation along the unit direction **v**_*k*_, scaled by the square root of its eigenvalue to match the natural magnitude of the latent space, and multiplied by an intervention scalar *σ* ∈ {−2.0, −1.0, 1.0, 2.0}:

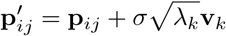

The patched representation **P**^′^ was then passed through the frozen distogram head to observe the mean change in entropy, 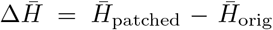. An asymmetric response—where positive *σ* sharpens the distogram (decreasing entropy) and negative *σ* blurs it (increasing entropy)—demonstrates causal control over geometric certainty. To ensure these effects are uniquely tied to the learned PC axes, identical shifts were performed using random unit vectors of matched magnitude 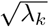 as a noise baseline.

#### 4.6.3 Transplanting Latent Certainty

Finally, to test whether geometric confidence is encoded in a universal, sequence-independent spatial language, we performed a cross-protein patching experiment. We extracted the pair representation from the highest-accuracy protein in the training set (the source, *S*) and transplanted it into low-accuracy, high-entropy test proteins (the targets, *T* ). For a target of length *L*_*T*_ and a source of length *L*_*S*_, the target’s representation was overwritten up to the pairwise minimum shared sequence length *L*_min_ = min(*L*_*T*_, *L*_*S*_):

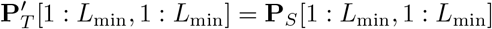

If cross-protein transplantation successfully induces a massive entropy collapse 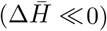 in an otherwise uncertain target protein, it empirically demonstrates that the network’s geometric certainty is an autonomous, transferable property of the latent space that transcends specific amino acid sequences.

### 4.7 Adversarial Mutational Pipeline

#### 4.7.1 Point Mutation Application

To systematically assess the representational origins of mutational invariance, we subjected all proteins to an adversarial sequence perturbation protocol adapted from [21]. For each target sequence, we randomly selected subsets of residues and replaced them with deleterious point mutations to create mutated sequences at predefined mutational loads of 10%, 20%, 40%, 70%, and 80%. Mutations were preferentially drawn from amino acid replacements known to disrupt secondary structure. These mutated sequences were then fed back into AlphaFold 3 as novel sequence inputs.

This perturbation procedure was executed under two distinct prediction regimes: an MSA-conditioned pipeline and a no-MSA pipeline. In the MSA-conditioned pipeline, the model reused the MSA of the unmutated sequence, which, as noted by [21], causes no major difference in structural invariance, as MSAs naturally incorporate homologous variation at the perturbed positions. In the no-MSA pipeline, evolutionary context was entirely withheld, with both the `pairedMsa` and `unpairedMsa` inputs to AlphaFold 3 being explicitly omitted.

To isolate the structural impact of mutations from the baseline effects of MSA deprivation, all TM-scores were calculated relative to the unmutated prediction from the corresponding condition. Specifically, mutated MSA-conditioned structures were compared to the unmutated MSA-conditioned baseline, and mutated no-MSA structures were compared to the unmutated no-MSA baseline. This matched-reference approach ensures that the measured structural divergence reflects true mutational sensitivity rather than a compound penalty for lacking co-evolutionary context.

Because the mutated and unmutated sequences for a given protein possess identical lengths *L*, their resulting latent tensors share the exact same dimensionality. This allows for direct comparison without the need for pooling or spatial aggregation. Let the single representation tensor be **S** ∈ ℝ^*L*×384^ and the pair representation tensor be **P** ∈ ℝ^*L*×*L*×128^. For these analyses, the single representation was flattened to a vector **s** ∈ ℝ^384*L*^, and the pair representation was flattened exclusively across its upper-triangular indices to a vector **p** ∈ ℝ^64*L*(*L*−1)^.

#### 4.7.2 Representation Drift and Uncertainty Calculations

Cosine distances were calculated directly on these full, unpooled vectors to compare the unmutated (**p**) and mutated (**p**^′^) states:

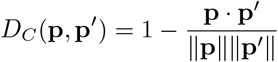

This formulation preserves position-specific representational drift without information loss. This parallel design allows us to isolate the degree to which representational robustness under extreme sequence drift is an autonomous biophysical property of the neural network versus a phenomenon reliant on MSA-derived homologous buffering.

Finally, representational uncertainty was quantified using the distogram entropy framework established in the previous section. Recall that the geometric uncertainty at each residue pair (*i, j*) is defined by its Shannon entropy, 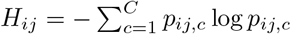. To yield a single global uncertainty metric 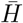 for the entire protein, these individual pairwise distributions were mean-pooled strictly across the upper triangle of the distogram:

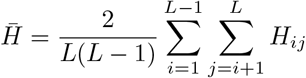

By computing 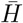 for both the MSA and no-MSA conditions, we establish a global counterpart to the sequence-separation Δ*H* analysis, allowing us to evaluate the network’s overall geometric certainty across the entire structural graph.

#### 4.7.3 Convergence of Drift Vectors in PCA Space

To quantify whether representational collapse under mutation follows a shared geometric trajectory rather than protein-specific failure patterns, we measured the directional coherence of drift vectors in PCA space. For each protein *i* at a given mutation level, we computed the drift vector 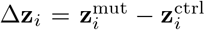, where 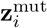 and 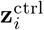 are the PCA projections of the mutated and unmutated pooled *C*_*N*_ pair representations, respectively. Each drift vector was normalized to unit length. To assess directional coherence across proteins, we computed the mean resultant vector length 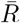, a standard measure of circular/directional concentration:

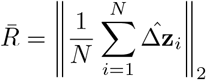

where 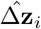 denotes the unit-normalized drift vector for protein *i* and *N* is the number of proteins. 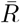 ranges from 0, indicating uniformly random drift directions, to 1, indicating perfect directional agreement across all proteins. Values approaching 1 therefore indicate that the representations collapse along a shared geometric pathway regardless of protein identity.

### 4.8 SABmark Alignment and Analyses

Comparing the internal representations of divergent proteins is challenging due to differing sequence lengths. To solve this without relying on spatial pooling—which smears and destroys position-specific geometric data—we utilized the gold-standard structural alignments from the SABmark benchmark.

For any given pair of proteins *A* and *B*, SABmark defines a subset of *K* structurally equivalent residue pairs. Let ℐ_*A*_ and ℐ_*B*_ represent the exact indices of these aligned residues for each protein.

#### 4.8.1 Unpooled Representational Similarity

Using this structural map, we extracted dimension-matched sub-tensors from the network’s full pair representations (**P**_*A*_ and **P**_*B*_). By indexing the tensors strictly at their structurally aligned positions, we can isolate the specific geometric features of interest:

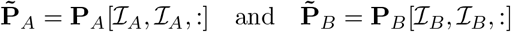

Because exactly *K* positions are extracted from both proteins, the resulting subtensors share an identical dimensionality of *K* × *K* × 128. This perfect symmetry allows us to fully flatten both sub-tensors into 1D vectors (**p**_*A*_ and **p**_*B*_) and directly compute their cosine distance, *D*_*C*_(**p**_*A*_, **p**_*B*_). By flattening rather than pooling, the spatial relationships between specific aligned residues are preserved in their entirety.

To rigorously test the significance of this representational similarity, we compared the results against two iteratively sampled baseline null distributions. First, a same-protein null was generated by computing the cosine distance between *K* random, strictly unaligned residues from the target protein pair. To ensure a stable baseline, this random sampling was repeated for 30 iterations per pair, and the resulting distances were averaged. Second, an unrelated monomer null was established to determine the expected similarity of completely random structural noise. For this, we drew two structurally distinct proteins from the 400 monomer dataset, compared *K* random residues between them, and averaged the distances over 20 random iterations.

#### 4.8.2 Structural RMSD via Kabsch Alignment

To evaluate whether this internal latent similarity translates to physical conformational accuracy, we calculated the Root Mean Square Deviation (RMSD) at the same *K* aligned positions using the Kabsch algorithm [60, 61].

Let **X**_*A*_ and **X**_*B*_ ∈ ℝ^*K*×3^ be the 3D coordinates for the alpha-carbon (*Cα*) atoms at the aligned indices. We first translate both sets of coordinates to their geometric center of mass, yielding 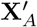 and 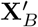:

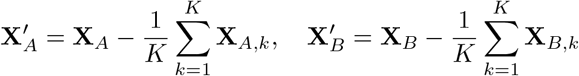

We then compute the covariance matrix 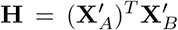 and perform Singular Value Decomposition such that **H** = **UΣV**^*T*^ . To ensure the resulting transformation is a proper rigid-body rotation and not a physical reflection, we apply a correction matrix **D** = diag(1, 1, sign(det(**VU**^*T*^ ))). The optimal rotation matrix **R** is calculated as:

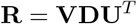

By applying this rotation, the structurally aligned RMSD between the two proteins is computed:

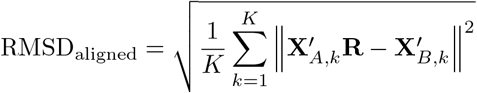

Parallel to the representational analysis, this aligned RMSD was evaluated against the same two null baselines (non-aligned residues and unrelated monomers) to ensure structural significance.

#### 4.8.3 Evaluating MSA Dependency

Both the representational cosine similarities and structural RMSDs were computed simultaneously for the standard pipeline (MSA) and the sequence-only pipeline (no-MSA). By analyzing the difference between these conditions (ΔRMSD = RMSD_no-MSA_ − RMSD_MSA_) specifically at the SABmark-aligned positions, we isolated the degree to which the network relies on evolutionary homology to correctly fold universally conserved geometric motifs.

#### 4.8.4 Nearest-Neighbor Fold Classification

To evaluate how well the internal representations support fold-level organization, we performed a nearest-neighbor classification over the SABmark protein set. For each protein, we computed the cosine similarity between its pooled pair representation and those of all other proteins in the set, using a leave-one-out scheme so that no protein is compared against itself. The nearest neighbor was identified as the protein with the highest cosine similarity, and the classification was scored correct if that neighbor belonged to the same SABmark fold group. Accuracy is reported as the fraction of proteins whose nearest neighbor shares their fold group. In the event of a tie, the neighbor was selected at random among the tied candidates, but ties were rare in practice and did not materially affect results.

### 4.9 Fold-Switching Protein Analysis

Because fold-switching is typically localized to a specific metamorphic subsequence, we first performed a windowed sequence alignment to identify the exact residue indices, denoted as ℐ_FS_, within the full predicted sequences of both conformations. To ensure comparative rigor, we also defined an internal control region, ℐ_ctrl_, of identical length *K* = |ℐ_FS_| for each pair, drawn randomly from the non-metamorphic, non-overlapping residues of the same proteins.

The degree to which the network’s internal geometric logic shifts to reflect these alternative states was quantified by extracting the Pairformer representations strictly at the metamorphic indices. Given the pair representations **P**_1_, **P**_2_ ∈ ℝ^*L*×*L*×128^, the extracted sub-tensors 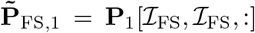 and 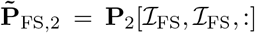 share an identical dimensionality of *K* × *K* × 128. We calculated the representational drift between these states using the unpooled cosine distance on the fully flattened vectors 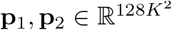.

Physical structural divergence at these sites was evaluated using the Kabsch algorithm to compute the RMSD at the ℐ_FS_ indices across three distinct comparative modes:

We first aligned the predicted metamorphic states to evaluate AlphaFold 3’s ability to distinguish alternative conformations for a single sequence. These predictions were then validated against their corresponding experimental PDB structures to ensure accuracy relative to the correct physical state. To isolate genuine signal from noise, results were compared against an internal baseline of non-switching ℐ_ctrl_ regions and an external null distribution. This null baseline was established by iteratively sampling random length-*K* regions from a diverse pool of unrelated proteins from the 400 monomer dataset, ensuring a stable estimate of stochastic representational similarity. Evaluating these metrics across both MSA and no-MSA conditions allowed us to isolate the network’s autonomous capacity to represent metamorphic states independent of evolutionary buffering.

### 4.10 MSA Perturbation Analysis

To systematically characterize how the Pairformer depends on different aspects of multiple sequence alignment information, we generate two classes of perturbed MSAs from the original inputs: column-shuffled and row-subsampled variants. These perturbations are applied to the unpaired MSAs in the AlphaFold 3 input JSON files. All paired MSAs are replaced with the query sequence, and all predictions are computed with identical hyperparameters and random seeds to the control.

#### 4.10.1 Column Shuffling: Destroying Co-evolutionary Coupling

Column shuffling permutes each alignment column independently across MSA rows, destroying the pairwise correlations between positions while preserving the marginal amino acid distribution at each position. The key algorithmic challenge is handling the A3M format, in which MSA rows contain insertions (lowercase characters) at different string positions relative to the query sequence [62].

An alignment column is indexed by position *i* among the query sequence’s aligned characters. Because insertions occupy different string offsets in different rows, alignment column *i* appears at a different string position in each row. The algorithm is:

##### Algorithm 1

Column Shuffling for MSAs

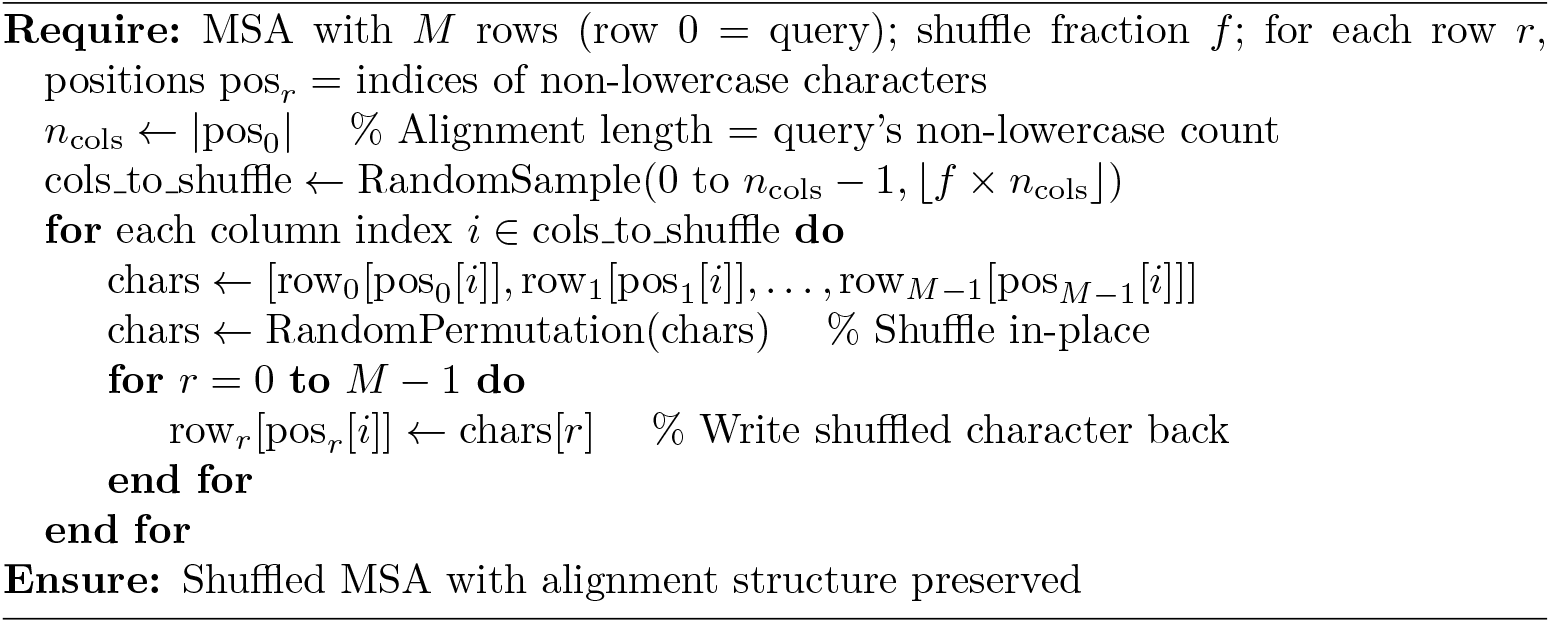

The critical property is that we permute characters within alignment column *i* across all *M* rows. Each row contributes one character and receives one character back, so the alignment length in each row is invariant. This preserves AlphaFold 3 featurizer compatibility [4]. We apply column shuffling at rates of 10%, 20%, 40%, 70%, 90%, 95%, and 99%.

#### 4.10.2 Row Subsampling: Reducing MSA Depth

Row subsampling randomly removes a specified fraction of homolog rows while keeping the query sequence fixed. The remaining rows retain their true pairwise co-evolutionary correlations but represent a shallower evolutionary context. We subsample at rates of 10%, 20%, 40%, 70%, 90%, 95%, and 99%. Homolog rows are selected uniformly at random without replacement. All subsampled MSAs are processed identically to the original.

#### 4.10.3 Phylogenetic Subsampling

To assess how phylogenetic character and MSA depth jointly determine structural accuracy, we constructed controlled MSA variants for 61 qualifying proteins selected from the 200 novel monomers. Qualifying proteins were required to have at least 10 sequences in each of three phylogenetic tiers: similar (≥ 80% identity to the query), medium (50–70% identity), and dissimilar (≤ 30% identity). Sequence identity was computed over aligned, non-gap positions only, excluding lowercase insertion characters from both numerator and denominator, consistent with the A3M alignment format used by AlphaFold 3 [4]. A fourth tier, random, was constructed by sampling uniformly from all homolog rows regardless of identity, serving as a matched-depth baseline with natural phylogenetic diversity. For each tier, subsets of 1, 5, and 10 sequences were generated. Within the similar tier, sequences were sorted by descending identity to the query and the top *n* were selected; within the dissimilar tier, sequences were sorted by ascending identity and the bottom *n* selected; within the medium tier, sequences were sorted by proximity to the tier midpoint (60% identity) and the *n* most representative selected. This deterministic sort order ensures that smaller counts are always strict subsets of larger ones, so that differences across depths reflect depth alone rather than sequence composition. Random sequences were drawn without replacement. All other AlphaFold 3 inputs were held constant across conditions. Structural outputs were compared against the full-MSA control prediction for the same protein, with the no-MSA prediction serving as the lower baseline. Recall for contact prediction is computed as TP / (TP + FN) with respect to the full-MSA control prediction.

#### 4.10.4 Fake MSA Injection

To determine whether MSA utility depends on evolutionary relevance to the query sequence or merely on alignment format and depth, we constructed fake MSAs by injecting sequences drawn from the MSAs of the 139 non-qualifying proteins into the input features of each of the 61 qualifying proteins. For each qualifying target protein with alignment length *L*, donor sequences were extracted from non-qualifying protein MSAs and trimmed to exactly *L* aligned columns by retaining the first *L* non-lowercase (aligned) characters along with any lowercase insertion characters immediately following them, and padding with gap characters if the donor sequence contained fewer than *L* aligned columns. This procedure preserves the A3M alignment format required by AlphaFold 3 while ensuring the injected sequences have no evolutionary relationship to the target. Donors were sampled without replacement across proteins, with at most one sequence drawn per donor protein before cycling to a second pass. Injection depths of 1, 10, 20, 50, and 100 sequences were tested. As with phylogenetic subsampling, smaller counts are strict subsets of larger ones. All other AlphaFold 3 inputs were held constant, and structural outputs were compared against the same full-MSA control and no-MSA baselines used in the phylogenetic subsampling analysis. Recall for contact prediction is computed as TP / (TP + FN) with respect to the full-MSA control prediction.

### 4.11 Statistical Tests and Structure Scoring

All structural comparisons—including TM-score calculations—were performed using the *OpenStructure* package [37, 63]. Correlation analyses were quantified using both Pearson’s correlation coefficient (*r*), which measures linear association, and Spearman’s rank correlation coefficient (*ρ*), which captures monotonic relationships independent of linearity, as implemented in the Python *Scipy* package [64]. Unless otherwise noted, all statistical tests, correlations, and regression analyses reported are significant at *p <* 0.05.

## 5 Acknowledgments

This research was supported in part by grant GM-118039 from the Division of General Medical Sciences of the National Institutes of Health. We thank Jessica Forness for proofreading and polishing this manuscript.

